# Inversions Dominate Evolution in the Highly Admixed European Sardine (*Sardina pilchardus*)

**DOI:** 10.1101/2025.01.15.633256

**Authors:** SJ Sabatino, M. P Cabezas, P Pereira, S Garrido, A Múrias, M Carneiro, PJ Talhadas, B Louro, C Cox, AVM Canário, A Veríssimo

**Affiliations:** CIBIO-InBIO, Research Center in Biodiversity and Genetic Resources, University of Porto, Vairão 4485-661, Portugal; BIOPOLIS Program in Genomics, Biodiversity and Land Planning, CIBIO, Vairão 4485-661, Portugal; Centre of Molecular and Environmental Biology (CBMA) and ARNET—Aquatic Research Network, Department of Biology, University of Minho, Gualtar Campus, 4710-057 Braga, Portugal; Institute of Science and Innovation for Bio-Sustainability (IB-S), University of Minho, Gualtar Campus, 4710-057 Braga, Portugal; Portuguese Institute for Sea and Atmosphere (IPMA), Lisbon, Portugal; Marine and Environmental Sciences Centre (MARE), Faculty of Sciences, University of Lisbon, Portugal; Faculty of Sciences, University of Porto, Porto, Portugal; Algarve Centre for Marine Sciences (CCMAR/CIMAR), Gambelas Campus, University of Algarve, 8005-139 Faro, Portugal

## Abstract

Inversions can play key roles in the genetic architecture of adaptation, but the scale of their effects across different species remains poorly understood. Here, we use whole-genome sequencing to investigate the influence of inversions on the population genomics of the r-selected European sardine (*Sardina pilchardus*). Allele frequency differences from millions of SNPs across 34 populations spanning the species’ range were analyzed. Genomic scans identified several extreme outlier regions overlapping chromosome-scale inversions, collectively representing over half the genome. Our findings suggest these inversions are associated with locally adapted life history strategies. First, SNPs within outlier regions containing inversions exhibited striking allele frequency differences between Atlantic and Mediterranean sardines, which differ in key adaptive life history traits. In the Atlantic, inversion allele frequencies varied latitudinally, while in the Mediterranean, they shifted longitudinally, aligning with temperature and oceanographic features that influence sardine life history strategies. Moreover, adjacent populations in contrasting environments displayed pronounced allele frequency differences in inversions. These spatial patterns of allele frequencies sharply contrasted with those based on neutral loci, indicating they are driven by selection. After rigorously filtering SNPs affected by selection and inversions, sardine populations showed high admixture across their range but significant population structure and isolation by distance, especially in the Mediterranean. This study demonstrates that inversions can shape genome-wide patterns of genetic diversity and population structure in highly admixed r-selected marine species. These findings also offer crucial insights for stock delimitation and management of this commercially valuable species in the face of climate change.

## Introduction

Structural variants, such as inversions, are widespread across the plant and animal kingdoms and can be important sources of genetic variation (Sturtevant, 1921; Dobzhansky, 1970). Inversions occur when a segment of DNA is excised from a chromosome and reinserted in the opposite direction, which often disrupts recombination in that region. This disruption of recombination stops or slows the segregation of mutations within inversions, allowing them to form haplotypes maintained through genetic linkage (Berdan et al., 2023; Charlesworth & Barton, 2018). While larger haplotypes targeted by natural selection would typically be broken down by recombination over a few generations (Berdan et al., 2023), within inversions, locally adapted, multi-gene haplotypes can evolve and persist. However, as beneficial mutations accumulate within adaptive inversion haplotypes, so can deleterious mutations that would normally be eliminated by recombination and selection or drift (Hill & Robertson, 1966). In extreme cases, the genetic load and meiotic incompatibilities associated with inversion haplotypes can be lethal or cause hybrid sterility, reinforcing the boundaries between diverging lineages (Jay et al., 2021). In less severe cases, inversions may segregate as balanced polymorphisms maintained by spatially or temporally varying selection, overdominance, or frequency-dependent selection (Haldane, 1957; Lewontin, 1964), among other forces. Due to their potential to affect fitness in multi-locus traits more significantly than single mutations, inversions can serve as key sources of adaptive genetic diversity across a range of ecological and evolutionary contexts.

Inversions may be especially important in species with specific genomic characteristics (Jay et al., 2021; Kirkpatrick, 2010) and ecologies. For example, in r-selected species with high gene flow that live in variable environments, recombination may make inversions beneficial for adaptive changes in quantitative traits (Berdan et al., 2023). Indeed, inversions were discovered in such species, *Drosophila melanogaster* and *D. simulans* (Sturtevant, 1921), and have since been shown to be important in their evolution (Nunez et al., 2024). Similar observations have been made in several others, including silversides (Akopyan et al. 2022), sticklebacks (Roesti et al., 2015), anchovies (Meyer et al. 2024), snails (Morales et al., 2019), crabs (Tepolt et al., 2022), and deer mice (Hager et al., 2021). Inversions in the Atlantic herring (*Clupea harengus*), which share some life history traits with the European sardine (*Sardina pilchardus*) studied here, are particularly relevant. In Baltic Sea herring, large-scale inversions exhibit sharp allele frequency differences among populations despite ongoing gene flow and control traits such as spawning time (Atmore et al., 2022; Jamsandekar et al., 2024). These adaptations allow herring with distinct life-history strategies to exploit alternative niches in the same locations and adapt to environmental gradients across their range. While the role of inversions in adaptation is well-documented, the ecological and evolutionary circumstances that determine the scale of their effects across species remain poorly understood.

In this study, we explore the population genomics of the European sardine (*Sardina pilchardus*), and how inversions have shaped it. This small pelagic fish is a highly fecund, r-selected species with extended pelagic egg and larval phases (lasting weeks) and adult, density-dependent, movement to find prey (Garrido et al., 2016; Silva et al. 2019). The species’ range covers the North and Central Atlantic Ocean, with populations extending from the North Sea to Senegal, as well as smaller groups in the Azores, Madeira, and Canary Islands (Parrish et al., 1989; Grant & Bowen, 1998), and across the Mediterranean and Black Seas. Consequently, sardines form a metapopulation of billions of individuals. One might, therefore, expect low or negligible population structure within the species, typical of many similar fish species (Palumbi et al. 2003). While some studies suggest that the European sardine is panmictic, others report significant genetic structuring (reviewed in Caballero-Huertas et al., 2022; da Fonseca et al., 2024), including an unexpectedly close genetic relationship between populations in the Eastern Mediterranean and those in the Canary Islands and Morocco (da Fonseca et al., 2024). Additionally, sardines exhibit considerable phenotypic variation in life-history traits, such as growth rate, size at maturity, and number of gill rakers (e.g., Silva et al., 2008; Dimarchopoulou & Tsikliras, 2022; Stratoudakis et al., 2007, 2008; Ganias et al., 2007; Baldés et al., 2022; Costalago et al., 2015). These traits distinguish two formerly named/recognized European sardine subspecies—*S. pilchardus pilchardus* in the Atlantic and *S. pilchardus sardina* in the Mediterranean — and vary latitudinally in the Atlantic (Regan, 1916; Andreu, 1969; Parrish et al., 1989). The genetic basis for these phenotypic differences remains unclear, with genetic and phenotypic plasticity potentially contributing to the variation. Four chromosome-scale inversions have been identified in the individual used for the current European sardine reference genome (GCA_963854175.1; Brittain, 2024; O’Leary et al., 2016). However, their role in shaping population genomics and phenotypic diversity has not yet been explored using a chromosome-scale genome assembly.

Here, we demonstrate that unexpectedly strong genetic differentiation among neighboring sardine populations and close relationships among geographically distant ones are primarily driven by nine chromosome-scale inversions. The spatial patterns of allele frequencies we revealed suggest that genomic regions containing chromosome-scale inversions are under selection in this species and contribute to alternative life history strategies adapted to local, regional, and basin-wide ecological variation. Our study provides the first comprehensive analysis of the range-wide population structure and genetic diversity of European sardines, where the effect of inversions is explicitly accounted for, offering new insights into their population genomics. These findings have important implications for understanding the role of inversions in the evolution of highly migratory pelagic fish and for protecting the European sardine, which faces increasing threats from climate change (Louro et al., 2019; Longo et al. 2024).

## Materials and Methods

### Sampling, DNA Extraction, Library Preparation, and Sequencing

European sardines (n = 1,027) were collected from 33 locations across the species’ range (Table 1). Samples were obtained from 30 of the 33 locations in 2019 (spring: n = 15; fall: n = 15), with the remaining three taken in the spring of 2021 (Table 1) during annual research surveys conducted in the North and Central Atlantic using small pelagic trawls, except for samples from the Aegean, Ionian, and Adriatic Seas, which were obtained from local fishermen. All fish were immediately frozen and later processed in the laboratory. Given the variability in size at maturity throughout the specieś range and the marked differences between Atlantic and Mediterranean sardines (e.g., Dimarchopoulou & Tsikliras, 2022; Silva et al., 2006), juvenile and adult samples were obtained using location-specific cutoff values of size at 50% maturity (L50): 18 cm for Atlantic samples, and 16 cm for Mediterranean samples. The final sample set comprised 13 juvenile and 21 adult samples, totaling 34 (Table 1). Muscle tissue for DNA extraction was collected from each individual and stored at -80°C.

**Table 1.** Sampling locations, developmental stage, and sex ratio of the 34 European sardine samples analyzed in this study. “Map Num.” corresponds to the location markers shown in Figure 1. Abbreviations for each population, here called “Codes”, are referenced throughout the text. The developmental stage (Juvenile or Adult) of individuals in each sample, as defined in the Methods section, is listed under “Stage.” Additionally, the sex ratio is reported for each sample when available.

| Map Num. | Sampling area | Codes | Basin | Lat. | Long. | Stage | Sex (F/M) |
| --- | --- | --- | --- | --- | --- | --- | --- |
| 1 | South Morocco | SMO_J | Atlantic | 23.668 | -16.452 | Juv. | 0,5 |
| 2 | Central Morocco | CMO_A | Atlantic | 27.05 | -13.568 | Adu. | 0,6 |
| 3 | Canary Islands | CAN_J | Atlantic | 28.038 | -15.349 | Juv. | 0,5 |
| 4 | Canary Islands | CAN_A | Atlantic | 28.521 | -16.119 | Adu. | 0,2 |
| 5 | Central Morocco | CMO_J | Atlantic | 30.117 | -9.701 | Juv. | 0,8 |
| 6 | North Morocco | NMO_J | Atlantic | 34.766 | -6.695 | Juv. | 1,0 |
| 7 | North Morocco | NMO_A | Atlantic | 35.676 | -6.59 | Adu. | 0,1 |
| 8 | Cadiz | CAD3_A | Atlantic | 36.398 | -6.418 | Adu. | 0,32 |
| 9 | Cadiz | CAD1_J | Atlantic | 36.838 | -6.792 | Juv. | unk. |
| 10 | Cadiz | CAD2_J | Atlantic | 36.955 | -7.007 | Juv. | 0,72 |
| 11 | South Portugal | SPA_A | Atlantic | 37.011 | -8.992 | Adu. | 0,5 |
| 12 | South Portugal | SPA_J | Atlantic | 37.432 | -8.823 | Juv. | 0,4 |
| 13 | Central Portugal | CPL_A | Atlantic | 39.208 | -9.378 | Adu. | 0,4 |
| 14 | North Portugal | NPO_A | Atlantic | 40.859 | -8.759 | Adu. | 0,2 |
| 15 | North Spain | NSS_A | Atlantic | 42.73 | -9.06 | Adu. | 0,5 |
| 16 | North Spain | NSC_A | Atlantic | 43.43 | -8.45 | Adu. | 0,5 |
| 17 | North Spain | NSB_J | Atlantic | 43.44 | -2.64 | Juv. | 0,4 |
| 18 | North Spain | NSA_A | Atlantic | 43.596 | -6.103 | Adu. | 0,2 |
| 19 | North Spain | NSR_A | Atlantic | 43.75 | -7.14 | Adu. | 1,0 |
| 20 | Central France | CFR_A | Atlantic | 44.061 | -1.391 | Adu. | 0,8 |
| 21 | Central France | CFR_J | Atlantic | 45.466 | -1.31 | Juv. | 0,6 |
| 22 | North France | NFR_J | Atlantic | 46.054 | -2.056 | Juv. | 0,6 |
| 23 | North France | NFR_A | Atlantic | 46.054 | -2.056 | Adu. | 0,6 |
| 24 | English Channel | ENC_J | Atlantic | 50.115 | -3.419 | Juv. | 0,6 |
| 25 | English Channel | ENC_A | Atlantic | 50.175 | -4.585 | Adu. | 0,8 |
| 26 | Bristol Channel | BRC_J | Atlantic | 50.689 | -4.95 | Juv. | 0,7 |
| 27 | Morocco | MMO_A | Mediterranean | 35.642 | -5.072 | Adu. | 0,5 |
| 28 | Torreveja | TOR_A | Mediterranean | 38.05 | -0.523 | Adu. | 0,3 |
| 29 | Tarragona | TAR_A | Mediterranean | 41.074 | 1.386 | Adu. | 0,4 |
| 30 | Lyon Gulf | LYG_A | Mediterranean | 42.806 | 3.483 | Adu. | 0,6 |
| 31 | Adriatic Sea | ADR_A | Mediterranean | 45.201 | 13.016 | Adu. | 0,6 |
| 32 | Ionian Sea | ION_A | Mediterranean | 38.817 | 20.497 | Adu. | 0,1 |
| 33 | Aegean Sea | AEG_J | Mediterranean | 40.618 | 23.928 | Juv. | unk. |
| 34 | Aegean Sea | AEG_A | Mediterranean | 40.836 | 24.514 | Adu. | 0,9 |

Genomic DNA was extracted from all sardines using the DNeasy Blood & Tissue Kit (Qiagen), following the manufacturer’s instructions, including the RNase A treatment. The DNA quality was assessed by agarose gel electrophoresis and spectrophotometry (Nanodrop), and concentrations were measured using the Quant-iT Picogreen Kit (Invitrogen). For each sample collection (n = 34; Table 1), DNA from 25 individuals was pooled in equal amounts and quantified. Pool-seq library construction and whole-genome shotgun sequencing (NovaSeq6000, Illumina) were performed by Macrogen (Seoul, Republic of Korea), generating 2x150 bp paired-end reads from 350-bp insert libraries (TrueSeq DNA Nano, Illumina) with a minimum coverage of 60x based on a genome size of 1,100 Mb.

### Quality Control of Raw Reads, Mapping, and Filtering

The quality of raw reads for each pool was evaluated using FastQC v0.11.8 (Andrews, 2010) and summarized with MultiQC v1.12 (Ewels et al., 2016). Adapter sequences and low-quality reads were removed using Trimmomatic v0.39 (Bolger et al., 2014) with the following parameters: (ILLUMINACLIP:TruSeq3-PE-2.fa:2:30:10 LEADING:30 TRAILING:30 SLIDINGWINDOW:5:20 AVGQUAL:30 MINLEN:100), and the resulting filtered reads reassessed with FastQC v0.11.8 and MultiQC v1.12. Quality-trimmed reads were mapped to haplotype one of the phased *S. pilchardus* reference genome (GCA_963854175.1) using BWA MEM v0.7.17-r1188 (Li, 2013). Alignments were sorted using SAMtools v1.14 (Li et al., 2009), and read groups were added with GATK AddOrReplaceReadGroups (McKenna et al., 2010). PCR duplicates were identified and removed with Picard v2.21.2 MarkDuplicates (Broad Institute, 2019). The read mapping quality for each pool was obtained with QualiMap v2.2.1 (Okonechnikov et al., 2015).

### SNP Calling and Allele Frequency Estimates

To identify SNP loci, the bam files for each poolseq dataset were analyzed with FreeBayes v0.9.21 (Garrison & Marth, 2012). The FreeBayes parameters used included: p 50 (number of chromosomes per pool), pooled-discrete, min-coverage 10, min-repeat-entropy 1, min-mapping-quality 60, min-base-quality 20, use-mapping-quality, and max-coverage 100 per pool. The FreeBayes analysis was run using all 34 pools simultaneously to maximize the detection of rare alleles but was computationally intensive. FreeBayes was run with use-best-n-alleles 2 to reduce computational costs, and loci with more than two alleles were filtered using custom Python scripts. The resulting VCF file was filtered for loci with quality scores below 30, minor allele frequency (MAF) < 0.05, or those showing strand bias (“SAF > 0 & SAR > 0 & RPR > 1 & RPL > 1”) using bcftools (Danecek et al., 2021). SNPs in haplotypes were then left-aligned using the norm function. Loci within ten base pairs of insertions or deletions identified in the FreeBayes analysis were excluded using custom scripts. A sync file of allele frequencies was generated with Popoolation2 (Kofler et al., 2011) using default parameters for diploid organisms (50 chromosomes for n= 25), retaining only high-quality SNP loci in the final, filtered, VCF file.

### Genome Scans, Population Structure, and Genetic Diversity (F_ST_ and H_E_)

A key objective was to estimate population structure and genetic diversity in European sardines using neutral, unlinked loci. Initially, we used PCAdapt (Luu et al., 2016) and BayPass2.1 (Gautier 2015) to detect non-neutral and linked SNPs (outliers) in our dataset that were filtered for this analysis. PCAdapt identifies outlier loci by performing principal component analysis (PCA) on genotype data to detect genetic variation associated with population structure. Loci are marked as outliers if their allele frequencies deviate significantly from neutral expectations. As selection often affects SNPs linked to targeted sites through genetic hitchhiking (Lewontin & Krakauer, 1973), we applied a sliding-window approach with 20 consecutive SNP loci and a 10 SNP step size. The most likely population cluster number (k) was chosen based on the lowest value that explained the most variance before plateauing. For the full 34 pools, PCAdapt identified k = 2 (Figure S1), splitting Atlantic and Mediterranean populations. However, when working with a large number of samples (n = 34), k = 2 can be biased towards PC1 (Janes et al., 2017). Thus, we performed separate PCAdapt runs for the 26 Atlantic and 8 Mediterranean samples. Significant outlier windows (FDR-corrected p-value < 0.05) from all three PCAdapt runs were then pruned for redundancy and merged if within 50kb. All SNP loci in the set of merged outlier windows were filtered.

Next, we used BayPass to identify individual outlier SNPs from the PCAdapt- filtered dataset. BayPass measures covariance in allele frequencies across populations and simulates statistical cutoffs to detect those that deviate from neutrality. BayPass was run using a subset of SNPs randomly chosen from across the genome at least one kilobase apart, yielding about 50k SNPs (hereafter referred to as “LD-pruning”). BayPass was executed with default parameters, including the recommended initial read count (d0yij) set to 10 (1/5 the minimum pool size in chromosomes). The test statistic used in BayPass is x^T^x, an analog of F_ST_ that uses the covariance in allele frequencies among populations to help mitigate the confounding effects of hierarchical population structure when detecting outliers (Gunther and Coop 2013). Next, a null distribution of x^T^x values based on neutral expectations was simulated using the empirical data. The simulated x^T^x values were then compared to those based on the empirical data, and loci exceeding the top 99th percentile were filtered from the dataset (hereafter referred to as the “soft-filtered” dataset).

Pairwise F_ST_ estimates and expected heterozygosity (H_E_) were calculated using the soft-filtered dataset with computing.fstats in the Poolfstat R package (Gautier et al., 2022). Genetic relationships among the 34 populations were visualized using heatmaps and dendrograms. We also estimated pairwise F_ST_ for samples based only on SNPs within outlier regions identified by PCAdapt. While linkage in outlier regions associated with inversions is likely higher than normal, LD pruning was conducted like above (hereafter referred to as the “outlier-only” dataset) for consistency with the soft-filtered dataset above.

An initial comparison of pairwise F_ST_ using the soft-filtered and outlier-only datasets showed differences in magnitude but strikingly similar spatial patterns (Figure S2; Table S1). In addition, elevated F_ST_ values in these two datasets varied non-randomly among regions (i.e., were clumped; Figures S3 and S4; see methods below) and overlapped with known inversions (Table S2), suggesting that our soft-filtered dataset was not free from bias. To address this, we tested a second filtering approach, aiming to hard mask regions showing evidence of being outliers. This was achieved by analyzing population pairs (see Figure S5) from the soft-filtered dataset that exhibited atypically high F_ST_ overall, especially between neighboring populations (Table S1), and had highly heterogeneous F_ST_ across entire chromosomes. Based on these criteria, of all of the pairwise F_ST_ comparisons made for our 34 samples, four stood out among the rest and were further studied (CAN_A/NSB_J CAD1_A/TOR_A CAN_A/SMO_J ION_A/NSB_J). For each of the four, we generated Manhattan plots of pairwise F_ST_ estimates for 20-SNP windows with a 10-SNP step size using Poolfstat v.2.2 (Gautier et al., 2022). For comparative purposes, F_ST_-Manhattan plots were also generated for an expanded set of pairs of populations, including all adult and juvenile samples from the same location using the same approach (Figures S3 and S4). We then visually inspected the Manhattan plots for each chromosome to identify regions of elevated **F_ST_** (several-fold higher) relative to the rest of the genome that were continuous, and at least 500kb long (Figure S5). The resulting set of outlier regions was then fully masked from the SNP dataset. As with the “soft-filtered” dataset, the resulting SNP dataset was pruned for linkage to obtain a set of 50k SNPs, which were then filtered using BayPass. Like in the pairwise F_ST_ filter, the Manhattan plot of x^T^x statistics from the BayPass analysis was also visually inspected for regions of linked outlier loci, and the one identified (Figure S6) was similarly masked. The resulting set of SNPs (hereafter referred to as the “hard-filtered” dataset) was then used to measure pairwise F_ST_ and H_E_ using Poolfstat like above, with the additional step of calculating 95% confidence intervals for F_ST_ estimates using the block-jackknife method from the same software. Differences in H_E_ between adult and juvenile population pairs were then tested using a paired T-test.

### Isolation by Distance

To investigate spatial patterns of genetic differentiation and the relevance of lower (<0.10) F_ST_ estimates (Palumbi, 2003), a generalized least squares (GLS) regression was performed to test for isolation-by-distance (Yang, 2004) using pairwise F_ST_ estimates and least-cost distances among sample collections. Least-cost distances were constrained by the upper 1,200 m of water depth, which separates the Canary Islands from the mainland, and were estimated using the Marmap *v1.0.10* package. The GLS model was run using R 4.42 (R Core Team 2021), accounting for covariance structure via the corMLPE v1 package (Clarke et al. 2002). Confidence intervals (95%) for model predictions were obtained through 1,000 replicate simulations of normal distributions.

### Identification, Population Genetic Analysis, and GO Enrichment of Inversions

Four chromosome-scale inversions were identified during the sardine reference genome assembly based on a single individual, suggesting that additional inversions might be segregating within the species. This hypothesis was supported by our outlier analysis using PCAdapt, which revealed several chromosome-scale outlier regions that did not overlap the four inversions identified in the reference genome. To identify candidate inversions, we again visually inspected Manhattan plots of pairwise F_ST_ values from four population pairs in our genetic analysis (see above). Candidate inversions were defined as genomic regions larger than 500 kb with continuously elevated F_ST_ several-fold above the genome-wide average. This analysis identified five additional candidate inversions (Figure S5; Table S2), which were subsequently examined in detail.

To help verify the existence of candidate inversions, we sequenced the genome of an additional individual using Nanopore long-read technology, which is well-suited for detecting structural variants. The specimen sequenced was collected by local fishermen off Porto, Portugal, in 2018, euthanized, flash-frozen in liquid nitrogen, and stored at - 80°C. DNA was extracted from muscle tissue using a phenol-chloroform protocol to minimize shearing. The extracted DNA was cleaned using magnetic beads and rinsed with ethanol and a cleaning buffer. Sequencing was performed with the Oxford Nanopore MinION, following the manufacturer’s instructions, across six runs to maximize yield. Adapter sequences were removed using Porechop (https://github.com/rrwick/Porechop), and the reads were mapped to the sardine reference genome using minimap2 (Li, 2018) with the “map-ont” option. Structural variants were analyzed using npInv (Shao et al., 2018), which detects inversions based on main alignments and sub-alignments. The npInv analysis was run using default parameters, except the minimum inversion size was set to 100 kb to reduce false positives. Only inversions flagged as “PASS” in the VCF file and supported by more than one read were retained, with overlapping intervals merged into a final set of candidate inversions.

The npInv analysis identified 57 candidate inversions spanning 482,327,974 bases (Table S2). Although the total length of these predicted inversions is substantial, it is consistent with findings from the reference genome individual. This consistency may reflect high heterozygosity for inversions in the Atlantic populations from which the reference genome and the individual sequenced here were sampled (see Results). However, identifying the breakpoints of inversions using npInv with an alignment of moderate coverage (23x) is not expected to be highly accurate. Consequently, we used these results primarily to assess the five candidate regions identified through F_ST_ analysis rather than to discover novel inversions. Each of the five candidate regions (F_ST_) was supported by overlapping inversions in the npInv results (Table S2), with an average overlap of 71.7%. Although only four of the five inversions had overlaps of 80% or higher, and additional research is needed to confirm their existence and true breakpoints, we refer to them as inversions and studied them along with the four identified in the NCBI reference genome.

We estimated allele frequencies for the four known and five candidate inversions. First, differences in allele frequencies (ΔAF) were calculated for all loci in the SNP dataset before outlier filtering (∼752,000 SNPs) using Popoolations2 (Kofler et al., 2011). For each nine inversions analyzed, SNPs in the top 25th percentile of ΔAF were considered diagnostic of divergent inversion haplotypes. Allele frequencies were polarized across all populations to associate the reference allele with putative alternative haplotypes using custom Python scripts. Pie charts visualizing inversion haplotype frequencies were generated using custom R scripts. Linear models were then used to test correlations between allele frequencies and latitude in the Atlantic and longitude in the Mediterranean. For comparison, a similar number of putatively neutral SNPs from the hard-filtered dataset were also analyzed with a linear model, and an ANCOVA was used to test for differences between the slopes (rate of Δ in allele frequency with distance) based on inversion versus neutral loci.

Gene Ontology (GO) enrichment analysis was conducted to assess whether inversions or outlier regions contained genes linked to fitness and adaptation. Separate analyses were performed for outlier regions identified by PCAdapt, each of the nine inversions, and all inversions combined. GO terms were extracted from the reference genome annotation on NCBI, and enrichment was tested using the topGO v2.38.1 (Rahnenfuhrer 2019) package in R. The weight01Count algorithm was applied to filter results, and redundancy was reduced using Revigo (Supek et al., 2011).

## Results

### Pool-seq Data And Genome Scans for Outliers

We generated pool-seq data for 34 sample collections: 26 from the North and Central Atlantic and eight from the Mediterranean. The pools had a mean sequencing depth of 65x, ranging from 61x to 78x. Variant calling identified 25 million SNP loci, filtered for quality, minor allele frequency (MAF), and indels. A total of 752,000 SNPs were retained for downstream analyses.

We generated two sets of putatively neutral, unlinked SNPs, differing in whether outlier SNPs or entire outlier regions were masked (i.e., soft- and hard-filtered datasets, respectively). In the soft-filtered dataset, PCAdapt analysis yielded 178,420 windows (20-SNP window, 10 SNP steps), averaging 9,076 bp. Based on likelihood scree plots (Figure S1), we chose k = 2 to analyze all 34 populations, k = 3 for Atlantic-only, and k = 4 for Mediterranean-only populations. Although the best k was clear for the first two analyses, it was less definitive for the Mediterranean, where k = 3 was used after determining that k = 2 and k = 4 yielded similar results. Manhattan plots of FDR-corrected p-values for SNP windows across the genome are shown in Figure 1. Windows with p-values above the top 5^th^ percentile were considered outliers. The majority of outliers were found in the analysis of all 34 populations (30,103), followed by the Atlantic (6,998) and Mediterranean (1,783) analyses, covering 303,367,492 bp. Redundant windows were pruned and merged if they overlapped or were within 50 kb, reducing the count to 493 windows, totaling 361,369,992 bp. SNPs within these outlier regions were filtered out, reducing the SNP count from 7,186,560 to 1,758,209. After LD-pruning, we retained 54,816 SNPs, with 548 identified as outliers in our individual SNP BayPass analysis (Figure S6), resulting in a final “soft-filtered” dataset of 54,276 SNPs. The hard-filtered SNP set involved masking F_ST_ outlier regions totaling nearly 607 Mb, representing 54% of the 112 Mb European Sardine genome. After filtering, LD-pruning, and BayPass analysis, the final SNP set comprised 50,049 SNPs.

**Figure 1.**
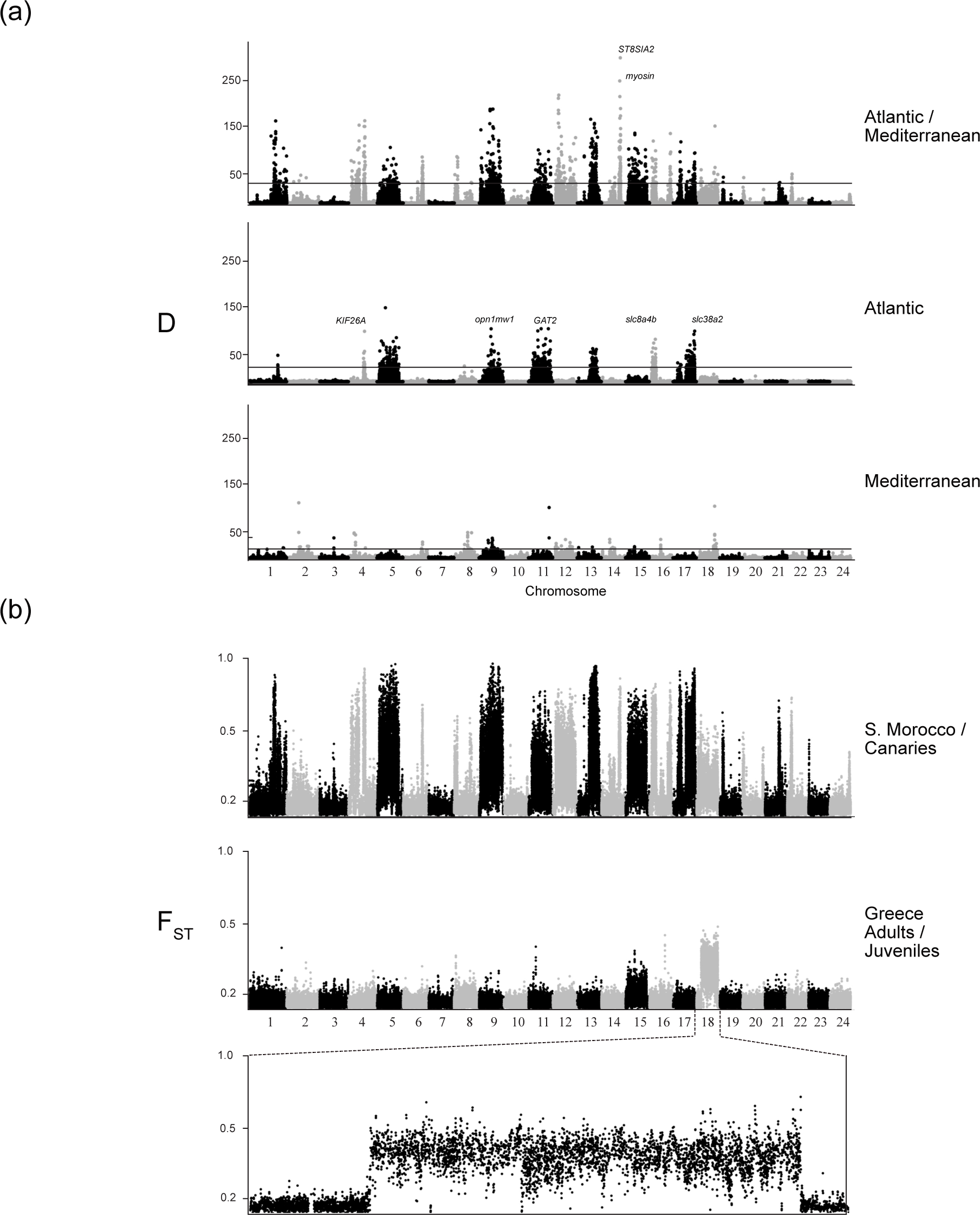
(a) Manhattan plots displaying p-values for Mahalanobis distances (DDD) calculated using PCAdapt, based on 20-SNP windows across the genomes of European sardine populations from different regions. The first plot includes all 34 populations in the study, the second focuses on the 26 Atlantic populations, and the third represents the eight Mediterranean populations. The Y-axis shows −log_10_ (p-values) for outlier detection, with the horizontal black line marking the 0.01 percentile threshold. (b) Manhattan plots of pairwise F_ST_ values calculated for 20-SNP windows (with a 10-SNP step) for two comparisons: Southern Morocco juveniles (SMO_J) versus Canary Islands juveniles (CAN_A), and Greek adults (AEG_A) versus juveniles (AEG_J).

### Genetic Diversity

Spatiotemporal patterns of genetic diversity were explored by estimating genome-wide averages of expected heterozygosity (H_E_) for each population. The H_E_ estimates we generated are correlated to the true H_E_, but given they are based on allele counts, they should be interpreted in a relative sense. The average and range of H_E_ for each filtered SNP dataset were as follows: soft-filtered 0.159 (0.156 - 0.161), hard-filtered 0.175 (0.160 - 0.180), and outlier-only 0.195 (0.170 - 0.20) (Table S3). Southern Iberian samples consistently had the highest heterozygosity, particularly near the Atlantic–Mediterranean transition zone, regardless of how SNPs were filtered dataset (Figure S7. This region aligns with the center of the species’ geographic range and the Atlantic–Mediterranean contact zone. In contrast, those populations at the southern, eastern, and to a lesser extent, northern edges of their sampled range had diminished H_E_. No significant differences in heterozygosity were found between adult and juvenile samples (paired t-test, p=0.423). However, H_E_ was higher in outlier regions when compared to the rest of the genome using the hard-filtered dataset (p<<0.05).

### Population Structure

We measured F_ST_ across the 34 samples of sardines to explore their genetic differentiation. Pairwise F_ST_ estimates varied significantly depending on the filtering method (Figure S2; Table S1). Maximum pairwise F_ST_ values ranged from 0.018 (hard-filtered) to 0.131 (soft-filtered) and averaged from 0.008 (Std. Dev. = 0.046) to 0.027 (Std. Dev. = 0.023), respectively. The maximum pairwise F_ST_ for the outlier-only dataset was 0.254, averaging 0.050 (Std. Dev. = 0.046). For the soft-filtered and outlier-only datasets, maximum F_ST_ values were several-fold larger than the average and standard deviation, which reflected the atypically strong divergence observed for the S. Morocco and Canary Islands populations compared to all others. However, all F_ST_ estimates, including those less than 0.05, were significantly different than zero (Figure S8), suggesting they accurately reflect the population dynamics of the species.

Clustering analysis with the soft-filtered dataset using Poolfstat identified seven spatially structured groups: 1) Biscay– English/Bristol Channel (U.K.), 2) NW Portugal– Galicia–C. Morocco, 3) S. Iberia, 4) W. Mediterranean, 5) E. Mediterranean, 6) Canary Islands, and 7) S. Morocco (Figure 2). The strongest differentiation occurred between S. Morocco, the Canary Islands, and all other populations. Atlantic samples differed from Mediterranean samples (average F_ST_ = 0.0942), with further subdivision within the Mediterranean between western and eastern basins (Figure 2). Additional breaks were identified in the Atlantic between Biscay–U.K. and NW Portugal–Galicia and between S. Iberia and other populations (Figure 2). Isolation-by-distance (IBD) patterns were evident in both the Atlantic (coefficient: 0.36, 95% CI: 0.33–0.39; Figure S9) and Mediterranean (coefficient: 0.17, 95% CI: 0.07–0.26), with exceptions such as a close relationship between Mediterranean and Canary Island populations, and the isolation of S. Morocco.

**Figure 2.**
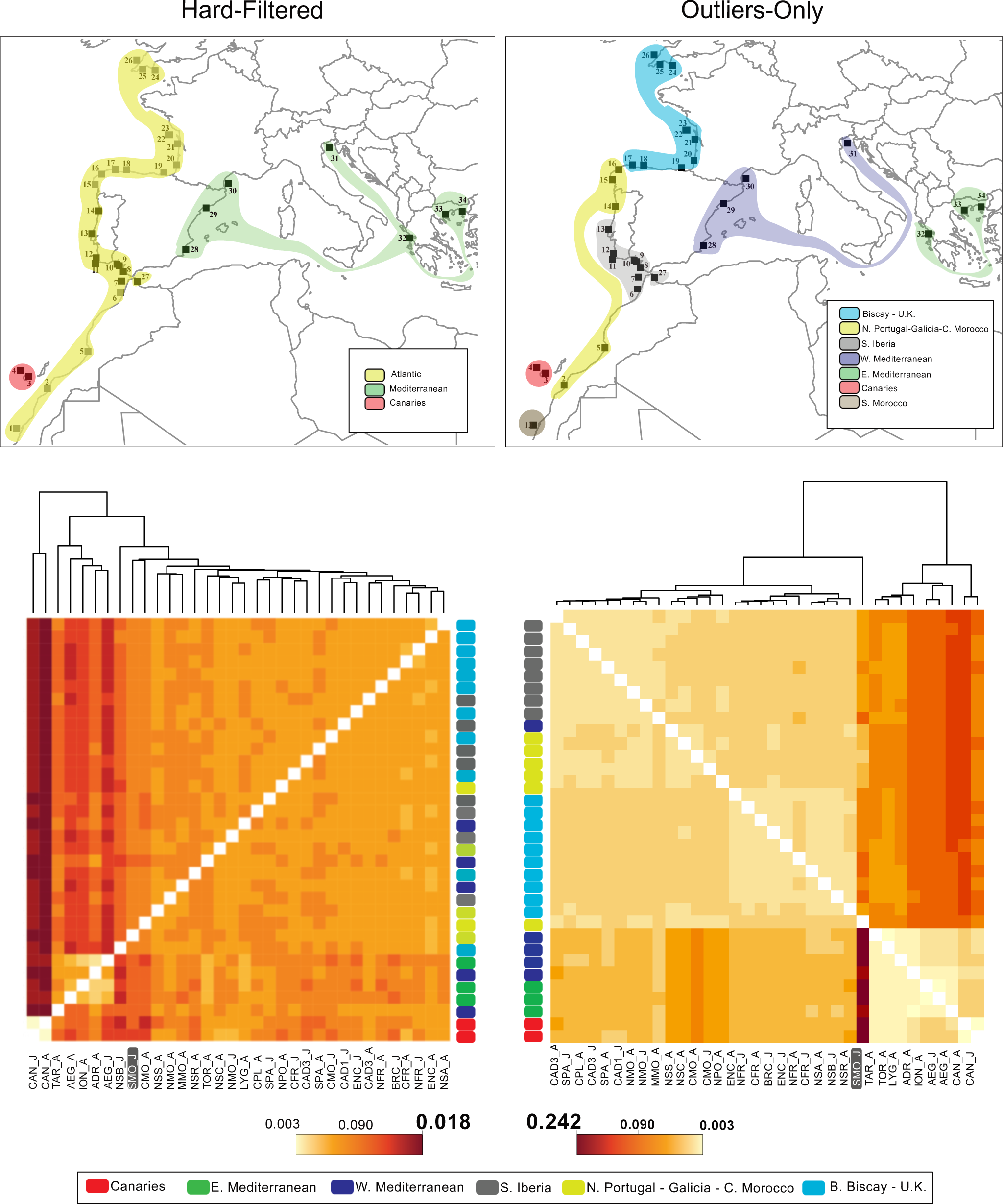
Maps depicting population clusters based on pairwise F_ST_ values, accompanied by heatmaps and dendrograms illustrating genetic relationships among populations. Results are shown for two datasets: the “Hard-Filtered” dataset, which excludes outlier loci associated with inversions and regions under selection (see Methods for filtering criteria), and the “Outliers-Only” dataset, which includes SNPs from loci likely influenced by inversions and selection. Numbers on the geographic maps correspond to sampling locations listed in Table 1.

In contrast, hard-filtered datasets revealed weak or no structure, with populations grouping into three main clusters: 1) Mediterranean, 2) Canary Islands, and 3) Atlantic (Figure 2). Relationships observed in the soft-filtered dataset, such as the Canary Islands’ connection to the Eastern Mediterranean, were not observed with hard filtering. Although F_ST_ values were consistently lower in the soft-filtered than the outlier-only datasets, the observed differentiation patterns in each were similar, suggesting residual biases due to inversions or selection in estimating gene flow. Therefore, we used the hard-filtered dataset to analyze migration, gene flow, and demographic history.

### Inversions: Spatial Patterns and GO Enrichment Analysis

We examined spatial patterns in allele frequencies of alternative haplotypes associated with known and candidate inversions in the European sardine to assess their impact on population genomics. Initially, we tested whether inversion haplotype frequencies correlated with the sex ratio in our samples. After adjusting for multiple comparisons, none of the inversions showed a significant correlation with sex.

We then analyzed spatial patterns in the allele frequencies of these inversions. Seven inversions (on chromosomes 5, 9, 11, 13, 15, 16, and 17) exhibited a consistent pattern, characterized by: 1) substantial differences between Atlantic and Mediterranean populations, 2) intermediate frequencies in most Atlantic populations, 3) similarity between the Canary Islands and the eastern Mediterranean, and 4) marked differences between Canary Island populations and those in Moroccan waters (Figure 3, Figure S10). Additionally, the S. Morocco population exhibited extreme divergence compared to all others. A second pattern, defined by inversions on chromosomes 12 and 18, showed similar frequencies across the sampling range, with moderate differences between Atlantic and Mediterranean populations. While most inversions displayed patterns deviating from neutral expectations (ANCOVA p<0.05; Figures 3, S9, S10, and S11), inversions on chromosomes 5, 11, and 17 exhibited the steepest spatial gradients relative to neutral expectations. In addition, inversions on chromosomes 15 and 18 showed significant differences between pairs of juvenile and adult samples from the same regions, suggesting potential changes in allele frequencies over short temporal scales or across generations.

**Figure 3.**
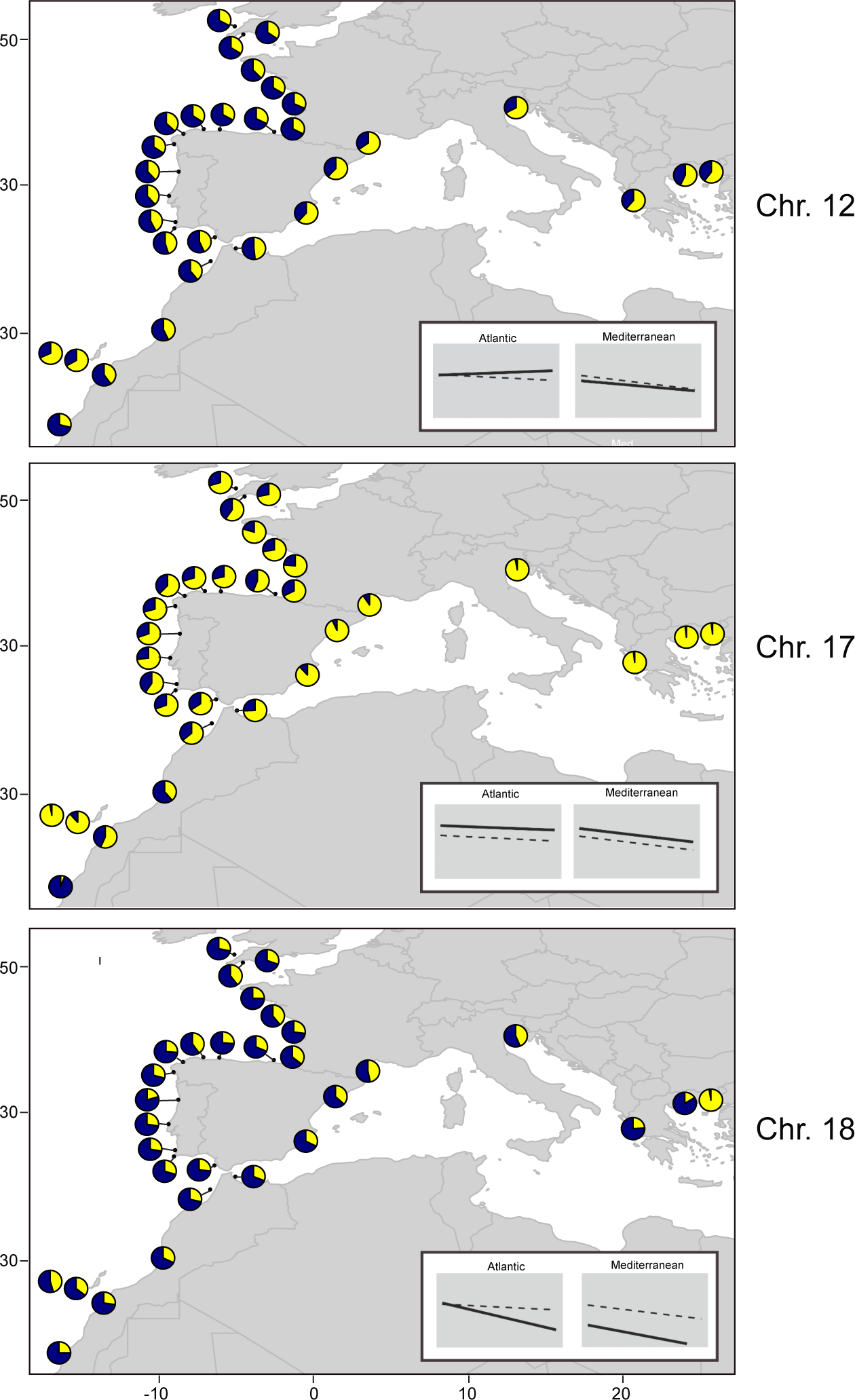
Estimated frequencies of alternative alleles (navy segments in pie charts) for chromosome-scale inversions identified in the European sardine populations. This figure highlights three of the nine analyzed inversions. Six inversions, exemplified by Chr. 17 (see Figure S11), show a shared pattern of relatively low alternative allele frequencies in the Canary Islands and Mediterranean populations compared to all others. In contrast, three inversions, including one on Chr. 12, exhibit alternative allele frequencies in the Mediterranean comparable to those in Atlantic populations. The inversion on Chr. 18, also shown here, illustrates how neighboring populations, such as those in Greece, can have strikingly different inversion allele frequencies, despite being otherwise highly admixed. Subplots within each map present trendlines comparing latitudinal (Atlantic) or longitudinal (Mediterranean) gradients in allele frequencies: dotted lines represent regressions for SNPs in non-inverted or outlier regions, while solid lines depict those for each inversion.

Lastly, we conducted Gene Ontology (GO) enrichment analysis on genes found within outlier regions and inversions to explore their association with known life history or fitness traits of sardines. In each set analyzed—outlier regions, individual inversions, and all inversions—genes were significantly enriched for various GO terms, many associated with fitness traits in sardines and other small pelagic fish (Table S4) (Berg et al., 2018; Miyazaki & Kobayashi, 2015, Sabatino et al. 2022b). Several of these genes that are related to brain development, eye function, movement (swimming), reproduction, and metabolism, such as *myosin*, *ST8 Alpha-N-Acetyl-Neuraminide Alpha-2,8-Sialyltransferase 2* (*ST8SIA2*), a member of glycosyltransferase family 29, *GABA transporter orthologue of the Solute Carrier Family 6* (*SLC6*), S*olute Carrier Family 8 Member 4b* (*slc8a4b*), a fish orthologue of the mammalian sodium/calcium transporter A1, *Solute Carrier Family 38 Member 2* (*slc38a2*), an amino acid/sodium transporter, and *Kinesin Family Member 26A* (*KIF26A*) were located at the peaks of inversion/outlier regions, suggesting they may be direct targets of selection in the European sardine.

## Discussion

Our findings provide new insights into the population genomics of the European sardine, revealing that up to nine chromosome-scale inversions, or regions of low recombination, have significantly shaped their population differentiation. These inversions appear to restrict gene flow in distinct parts of the genome, even amidst ongoing, high levels of admixture. As a result, we observe striking patterns of genetic differentiation at various spatiotemporal scales, including among adult and juvenile sardines within the same locations, between neighboring populations a few hundred kilometers apart, and across large water basins such as the North Eastern Atlantic and Mediterranean. The spatial and allele frequency patterns of these inversions revealed here, together with their correspondence to known patterns in life-history traits within the species discussed in detail below, suggest that they are under selection in the European sardine.

We also demonstrate that when outlier regions are fully masked, European sardine populations appear highly admixed throughout their range, with some restricted gene flow and isolation-by-distance (IBD). Our results offer the first comprehensive picture of population structure in the European sardine driven mainly by migration and genetic drift, without the confounding effects of selection and inversions. The observed mosaic of highly admixed versus strongly divergent genomic regions helps explain some conflicting results found in previous population genetic studies (Caballero-Huertas et al., 2022). These findings have important implications for managing one of Europe’s most economically and culturally significant fisheries species, especially regarding climate change.

### Genetic Variation For Inversions Correlates With Life History In Sardines

The spatial and allele frequency patterns for inversions in the European sardine shown here matched known variability in their life histories and the ecological factors thought to drive them. This includes the most consistent inversion allele frequency differences observed in our study, which were between Atlantic and Mediterranean sardines, as well as, latitudinal gradients in the Atlantic, and longitudinal ones in the Mediterranean. However, the Strait of Gibraltar, a well-known genetic barrier for many marine species, and re-current post-Pleistocene expansions from glacial refugia can generate patterns in allele frequencies similar to those we observed (Hewitt 2011). We show that these inversion patterns persist despite high levels of gene flow and strikingly contrast those based on neutral loci in non-inverted genomic regions, suggesting that they result from spatially varying natural selection. Concomitantly, widespread heterozygosity and intermediate inversion allele frequencies across their range imply that some form of balancing selection also shapes genetic diversity for these inversions. Here we discuss these findings and their implications for understanding the evolutionary history and population dynamics of the European sardine.

As seen in many fish species including salmonids, stickleback (Stearns 1992), and other clupeids, life-history traits in sardines evolve in response to ecological factors, resulting in locally adapted strategies (Neves et al., 2021). Traits such as growth rate, age or length at maturity, fecundity, egg size/quality, and feeding strategy (including gill raker number) are correlated in sardines (Brosset et al. 2016; Beauvieux et al. 2022; Costalago et al. 2015), with trade-offs often accompanying adaptations in specific traits (e.g., Ganias 2009). These trait correlations, combined with environmental gradients and regional differences, drive spatial and temporal variations in locally adapted life history strategies in sardines.

*Across Basins*—One of the clearest examples of life history variation that matched our results is for the only two named subspecies of sardines. Atlantic sardines are adapted to colder, nutrient-rich upwelling regions, such as off Morocco and Iberia. In these regions, sardines grow more slowly, mature earlier (da Fonseca et al., 2024; Silva et al. 2006, and 2008), and have more gill rakers (except in Azores, Madeira, and Canary Is.), a specialization for passive filter-feeding (low-energy investment) on microplankton (Andreu 1969; Garrido et al. 2007; Costalago et al. 2015). Mediterranean sardines inhabit warmer, less productive waters, reaching smaller total lengths and L50 than Atlantic sardines (Silva 2006). They also exhibit fewer gill rakers (similar to those in Atlantic islands) adopting a feeding strategy that almost exclusively relies on foraging via particulate-filter feeding on mesozooplankton, and thus consuming a significantly lower proportion of microplankton compared to Atlantic sardines (Andreu 1969; Garrido et al. 2007; Garrido and van der Lingen 2004, Chia-Ting et al., 2021). This flexible feeding behavior enables Mediterranean sardines to thrive in environments with lower and more variable plankton abundance (Bachiller et al., 2020; Ganias 2009).

*Within Basins*—The fitness traits that broadly distinguish Atlantic and Mediterranean sardines also vary within these basins in response to local environmental factors. For instance, peak spawning occurs at different temperatures in the Atlantic, ranging from 14.0–15.0°C in the English Channel to Western Iberia and 16.0–18.0°C in northwest Africa (Coombs et al., 2006; Bernal et al., 2007). These peak spawning temperatures could reflect species-wide tolerances (13.0–18.0°C) that coincide with local spikes in phytoplankton abundances or favorable oceanographic conditions for larval survival (Silva et al. 2006; Ferreira et al. 2023). However, subtle temperature changes within this range can negatively affect fitness (Garrido et al., 2016), supporting the hypothesis that sardines are locally adapted to specific thermal conditions. Further, sardines in upwelling regions like off Morocco and Iberia, where phytoplankton is abundant, have more gill rakers and specialize in filter-feeding (Andreu, 1969; Costalago et al., 2015). In contrast, populations in regions with lower or more unpredictable productivity, such as the North Atlantic and Canary Islands, have fewer gill rakers (Andreu, 1969; Bachiller et al., 2020). Growth and maturity patterns also vary. Sardines in the Bay of Biscay grow rapidly during short, warm summers to enhance overwinter survival, delaying sexual maturity. In colder regions with longer growing seasons, such as Iberia and Morocco, they grow more slowly and mature earlier (Silva et al., 2006, 2008). These patterns support the hypothesis of four broad environmental regions driving sardine reproductive cycles and life-history evolution—Biscay Bay, Portugal-Morocco, Western Sahara- Mauritania, and the Mediterranean (Coombs et al., 2006; Silva et al., 2008)— corresponding to the observed inversion allele frequency clusters (Figure 2), and spatial gradients in the Atlantic and Mediterranean (Figure S7).

*Within Regions—*Remarkably, the three populations studied that were, by several orders of magnitude, the most divergent for inversion alleles—the two from the Canary Islands and one from southern Morocco—fit these broader patterns of life-history adaptation. Sardines in the oligotrophic Canary Islands resemble Mediterranean sardines in traits such as gill raker number and length-at-age (Mata, 1997), reflecting adaptations to similar environmental conditions (Figure 2). Seven chromosome-scale inversions were more similar between Canary Island and Eastern Mediterranean sardines than their immediate neighbors, providing striking evidence for local adaptation and possibly parallel evolution (Häkli et al. 2017). Similarly, the southern Moroccan population (SMO_J) was the most southerly we sampled and is from an area with colder temperatures and more continuous and intense upwelling than found in neighboring areas, making it one of the most productive fisheries on the planet (Sarre et al. 2024). High frequencies of nearly all nine inversions had pervasive impacts on the genomes of individuals in the SMO_J sample, suggesting they are involved in adaptation to their unique environment. The striking contrast between populations in southern Morocco and the Canaries is as extreme as those between the Mediterranean and Atlantic subspecies, suggesting that this region is a microcosm of life history evolution that defines the species.

*Across time and space*—An alternative, or complimentary, hypothesis is that these sardine inversions segregate between distinct ecomorphs living in sympatry, as seen in the Atlantic herring (Bekkevold et al., 2007) and anchovy (Catanese et al., 2017). Like in herring, a potential candidate trait differentiating sardine ecomorphs is spawning time. Sardines exhibit variable spawning seasons across their range, influenced by fish size, age, and local thermal regimes (Stratoudakis et al., 2007). Another possibility is deep versus shallow water morphs that exploit different niches like in the anchovy (Catanese et al., 2017), perhaps to minimize competition (Gomes et al. 2001). However, distinct ecomorphs have not been widely reported in sardines, and intermediate inversion allele frequencies across most Atlantic populations suggest high levels of admixture rather than stable divergence. Future research should investigate the potential co-existence of ecomorphs, particularly in transition zones between life history types such as the Strait of Gibraltar, and assess correlations between inversion haplotypes and life-history traits.

Interestingly, the GO terms enriched within inversions and the traits linked to population differentiation are closely related, further supporting our findings. Specifically, brain development, eye function, and movement (swimming) are central to larval/juvenile development, feeding activity, reproduction, metabolism, growth, and maturity. Although our results suggest natural selection influences inversion allele distributions, genome-wide association studies (GWAS) and experimental validation of allele fitness are needed to rigorously test this hypothesis (Storz, 2005). However, GWAS for (quantitative) life-history traits is often challenging, especially in regions of low recombination like those seen here, where hitchhiking effects obscure selection signals. Comparative genomics and methods such as the McDonald-Kreitman test may provide a pathway to identify specific genes under selection within these inversions.

### Genomic Insights for Managing European Sardine Populations

Our study indicates that the European sardine is highly admixed throughout this range, consistent with its life history and known population size. However, the scale of population differentiation among sardine populations was highly dependent on how stringently outliers and inversions were filtered from the genome. We found pronounced spatial genetic heterogeneity along the sampled range using our soft-filtered dataset (Figure 1; Figure S2), which matched patterns found using the outliers-only dataset (Figure 2). In contrast, weak but significant genetic structure and IBD when outlier regions were fully masked (i.e., using the hard-filtered dataset). It is possible that the filtering methods used to generate the hard-filtered dataset were too aggressive. However, including even tens of outlier SNPs in an otherwise neutral dataset generates patterns of population structure produced by inversions (Figure S4). This sensitivity stems from genetic differentiation due to migration and drift alone, likely being weak, allowing for even small fractions of outlier loci in a dataset to spike F_ST_ estimates.

Further support for our filtering choices and related results comes from otolith studies on population structure in sardines. Restricted gene flow, admixture among neighboring populations, and population structure coinciding with areas of upwelling and phytoplankton production are supported by otolith studies (Correia et al. 2014; Neves et al., 2021). These studies also provide evidence of widespread dispersal and gene flow, including across the Strait of Gibraltar, but especially among populations in the Atlantic. These findings concord with our results for the hard-filtered dataset, suggesting they reflect accurate estimates of population differentiation and diversity.

The results from our study help reconcile some findings in previous studies that were difficult to understand. Genetic and genomic studies aiming to delimit the management/biological units in the European sardine have used a wide variety of genetic markers and approaches (i.e., allozymes, mtDNA, microsatellites, RADseq, and SNPs), often yielded conflicting results (Gonzalez & Zardoya, 2007; Kasapidis et al., 2012; Baibai et al., 2012; McKeown et al., 2024; reviewed in Caballero-Huertas et al., 2022). However, as previously suggested by several authors, we find that these discordances can be reconciled considering the component of the genome sampled in each case, i.e., neutral and non-neutral.

For instance, genetic homogeneity and weak IBD were reported along large spatial scales in studies using nuclear (neutral) microsatellites, mitochondrial DNA, or RADseq-derived SNPs (Gonzalez & Zardoya, 2007; Kasapidis et al., 2012; Baibai et al., 2012; McKeown et al., 2024; da Fonseca et al., 2024; Antoniou et al., 2023). In contrast, pronounced spatial genetic heterogeneity was detected previously using non-neutral allozyme markers (Ramon & Castro, 1997; Chlaida et al., 2006, 2008; Laurent et al., 2007) or SNP markers across regions of marked environmental shifts (Antoniou et al., 2023; da Fonseca et al., 2024). In short, the previously unknown scale of the impact of the inversions on the population genomics of the European sardine may have led to the major discrepancy in the patterns of population structure among studies. Only by conservatively considering outliers and inversion, which impact more than half the genome, can range-wide patterns of population structure and diversity be revealed.

Our observation of rapid inversion allele frequency shifts between adult and juvenile sardines in the North Aegean Sea and central Morocco (Figure S4) raises the possibility that selection in early life stages is very strong or that genetic drift plays a significant role over short time scales. Given that selection on larvae and juvenile fish is strong (Perez & Munch (2010) and the expected effective population sizes of sardines are large (Laurent and Planes 2007), the effects of selection may closely track seasonal or yearly environmental fluctuations, as seen in other r-selected species (Bitter et al. 2024). Supposing selection is not responsible for differences in inversion allele frequencies between adults and juveniles, our findings highlight how sharply drift can act in this species, probably due to high variance in reproductive success. Consequently, even moderate changes in connectivity or census size may have significant negative impacts on population differentiation and genetic diversity, despite their large meta-population effective size.

### Implications for fisheries management

Our findings on the population structure of European sardines throughout its range indicate a mismatch between the locally adapted genetic stocks detected here and the stock units currently defined for fisheries stock assessment and management. In European Atlantic waters, three sardine stocks are considered - a Northern stock (Celtic Sea and the English Channel), a Central stock (Bay of Biscay), and a Southern stock (Cantabrian Sea to the Gulf of Cadiz) (ICES, 2016, 2019), while three other stocks are assumed for Atlantic Morocco - a smaller stock off northern Morocco (between 32°N and 35°45′N); a large central stock (between 26°N and 32°N); and a southern stock from the south boundary of the species range northwards to 26°N. Here, we found no differences between the Northern and Central stocks, which instead formed a single genetic cluster extending into the Cantabrian Sea. The Southern stock was also shown to include different genetic clusters, notably one in NW Iberia and another one in southern Iberia extending into northern Moroccan waters and the adjacent Mediterranean Sea. Within the Mediterranean Sea, the General Fisheries Commission for the Mediterranean has established thirty management areas based on political and statistical considerations. However, we were only able to distinguish western vs. eastern genetic clusters. Our study highlights the need to further assess stock structure and dynamics in the European sardine, calling for international cooperation beyond European borders to update stock assessment and management strategies of this important shared fishery resource.

Our findings also underscore the importance of assessing and integrating spatial and temporal variability in European sardines’ population structure and diversity in future management plans and spatial patterns in the species’ location adaptation. Importantly, current boundaries among adjacent genetic stocks may vary annually depending on local environmental conditions and source-sink dynamics (Silva et al. 2019). The maintenance of all population units and their unique genetic diversity may be crucial for maintaining critical levels of biocomplexity to sustain fisheries at levels seen over the past few hundred years (Hilborn et al., 2003; Ruzzante et al., 2006; Kess et al., 2019; Sabatino et al. 2022a). Given how many life history traits in the species are tied to water temperature, much of the adaptive genetic variation we identified here will be critical for sardine populations’ long-term survival and stability as climate change (Sakamoto et al., 2022) continues to alter ocean temperatures and conditions.

## Author Contributions

Conceptualization: SJS, AV, AM; Funding: AV, AMS, SJS, AVMC, BL, and CC; Writing original draft: SJS; Review and editing: All authors, especially AV; Field Sampling: AV, SJS, MPC, and SG; Lab Work: MPC with contributions from SJS, PP, and AV; Analysis: SJS with contributions from MPC, PP, and AV.

## Acknowledgments

This work was performed under the scope of Project SARDINOMICS: Desenvolvimento de ferramentas moleculares para o melhoramento do conhecimento e da gestão dos stocks de Sardinha (PO Mar2020, MAR-01.04.02-FEAMP-0024) financed by the European Maritime and Fisheries Fund (EMFF), through the MAR2020 Operational Programme. Additionally, BL, SC, and AVMC received Portuguese national funds from FCT - Foundation for Science and Technology through projects UIDB/04326/2021, UIDP/04326/2022 and 2023, and LA/P/0101/2020. We are indebted to many colleagues and institutes who have helped collect the samples needed for this study: Jeroen Van Der Kooij (CEFAS); Erwan Duhamel (IFREMER); Alba Jurado, Cristina Bulto, Isabel Riveiro, Pablo Carrera, Teresa Garcia Santa-Maria (IEO); Anabel Colmenero Ginés (ICM-CSIC); Malika Chlaida, Salaheddine Sbiba (INHR); Carlotta Mazzoldi (University of Padova); Ada Chatzispyrou (HCMR); Chrysoula Gubili (FRI & DEMETER); Evina Gontikaki (HFRI). We also thank members of Evolgen at CIBIO-InBIO and CCIMAR for their helpful discussions on the subject matter.

## Data Availability

All genetic data for this study will be made available on GenBank (https://www.ncbi.nlm.nih.gov/genbank/) following publication.

---

**Figure S1.**
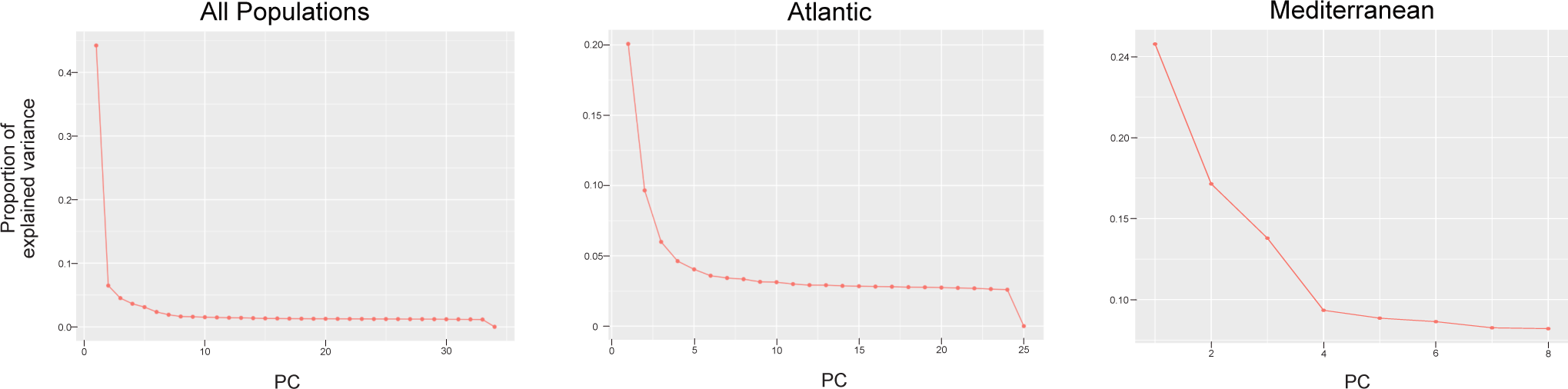
Scree plots from the three PCAdapt analyses: “All Populations” includes all 34 European sardine populations, while the “Atlantic” and “Mediterranean” analyses focus on populations from their respective regions. The optimal number of population clusters (KK) was identified as the smallest value where the proportion of variance explained plateaued, indicating diminishing returns with additional components.

**Figure S2.**
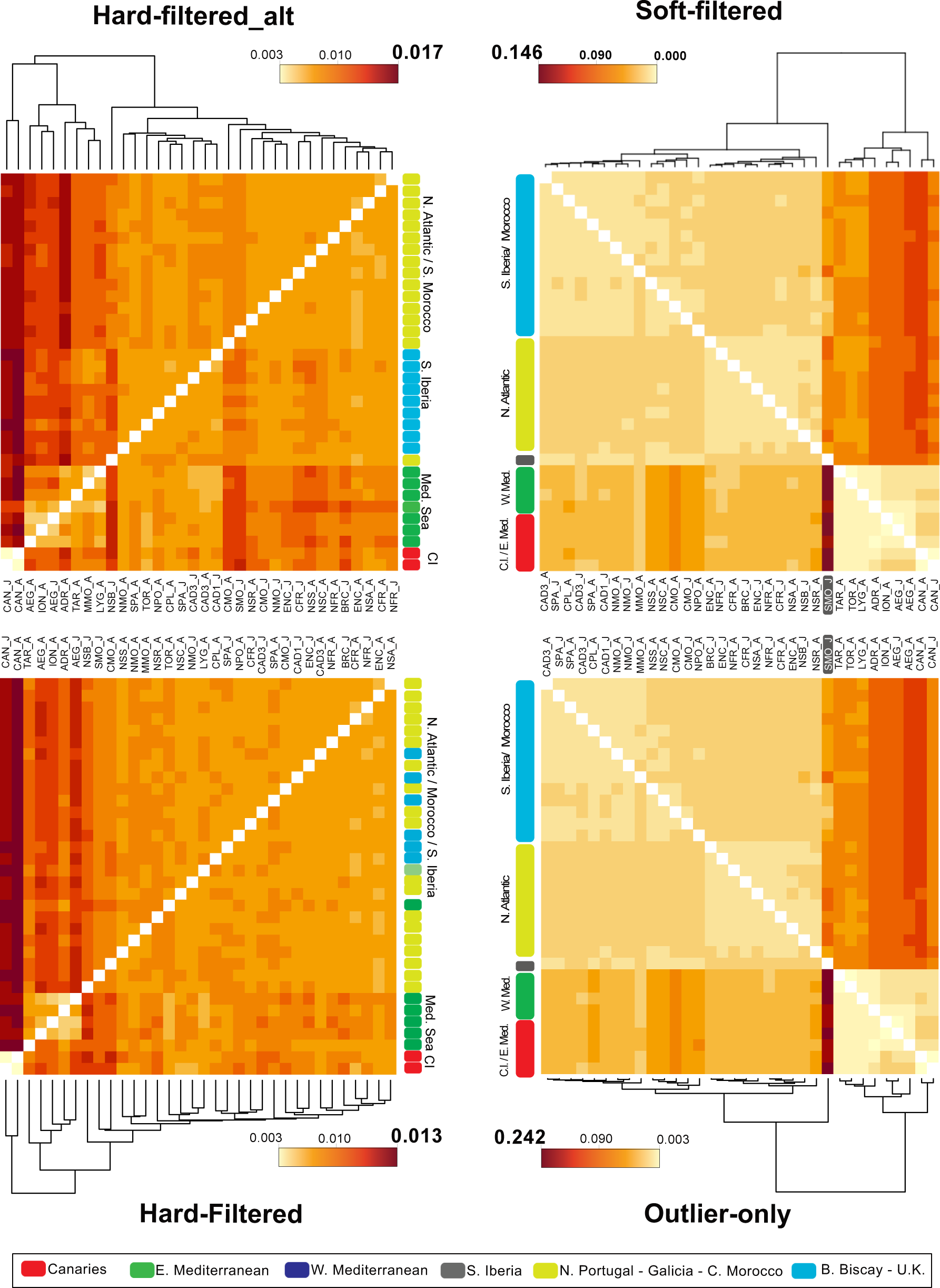
Genetic relationships among all 34 European sardine populations, represented as heatmaps and dendrograms based on pairwise F_ST_ values. Results are presented for four datasets: the “Soft-Filtered” dataset, which is filtered for outlier loci associated with inversions and selection; the “Hard-Filtered” dataset, which excludes all regions with outlier loci (see Methods for filtering criteria); the “Outliers-Only” dataset, which includes only SNPs from loci likely influenced by inversions and selection; and the “Hard-Filtered_alt” dataset, a variation of the “Hard-Filtered” dataset that includes a single outlier region identified in the final step of the BayPass analysis, demonstrating the dataset’s sensitivity to outlier inclusion.

**Figure S3.**
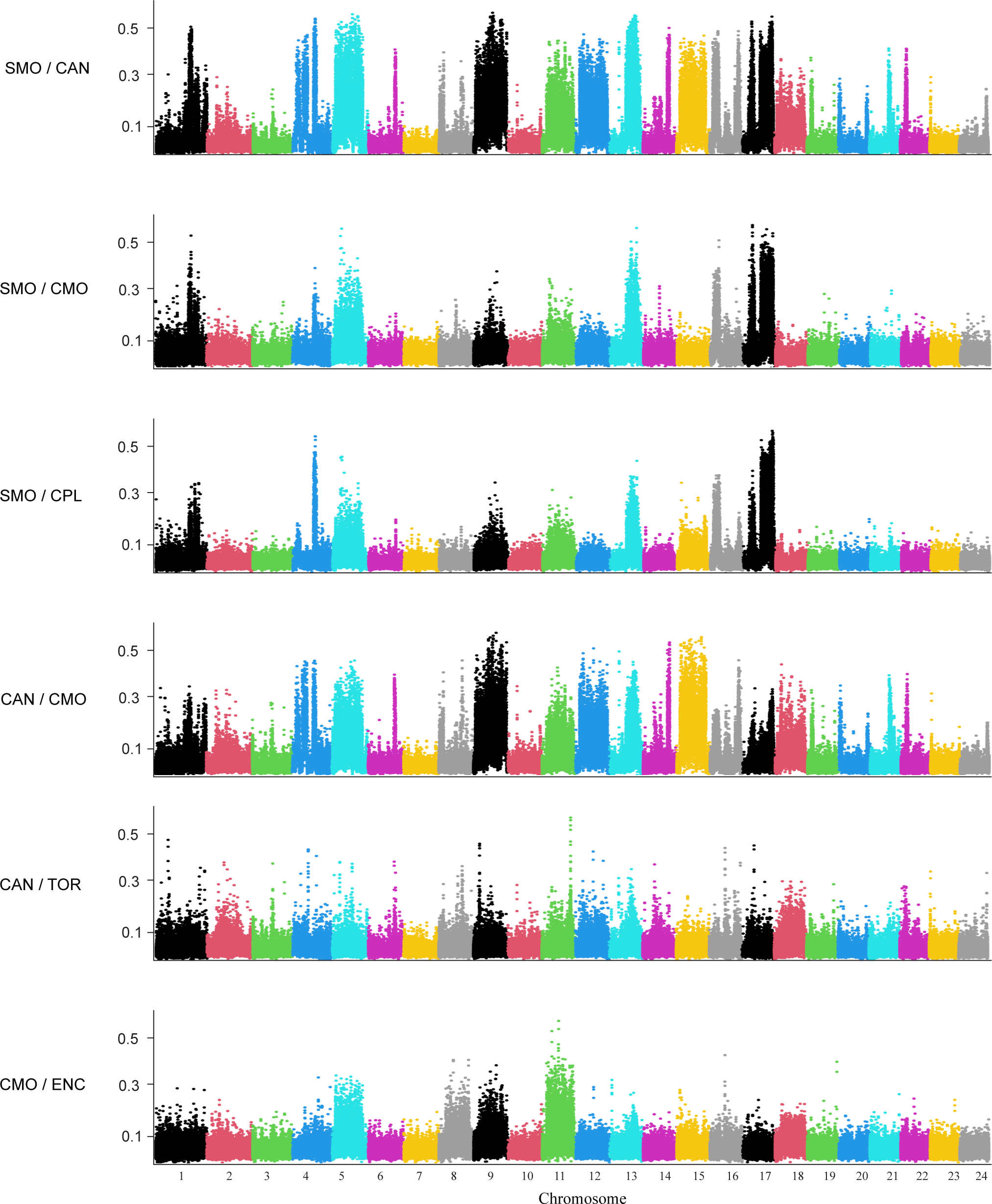
Pairwise F_ST_ values for 20-SNP genomic windows for selected population pairs that deviate from the typical Isolation by Distance (IBD) pattern in the European Sardine. Population codes are as follows: SMO_J (Southern Morocco), CAN_J (Canary Islands), CPL_A (Central Portugal),TOR_A (Western Mediterranean), and ENC_A (English Channel) (see Table 1 for more details).

**Figure S4.**
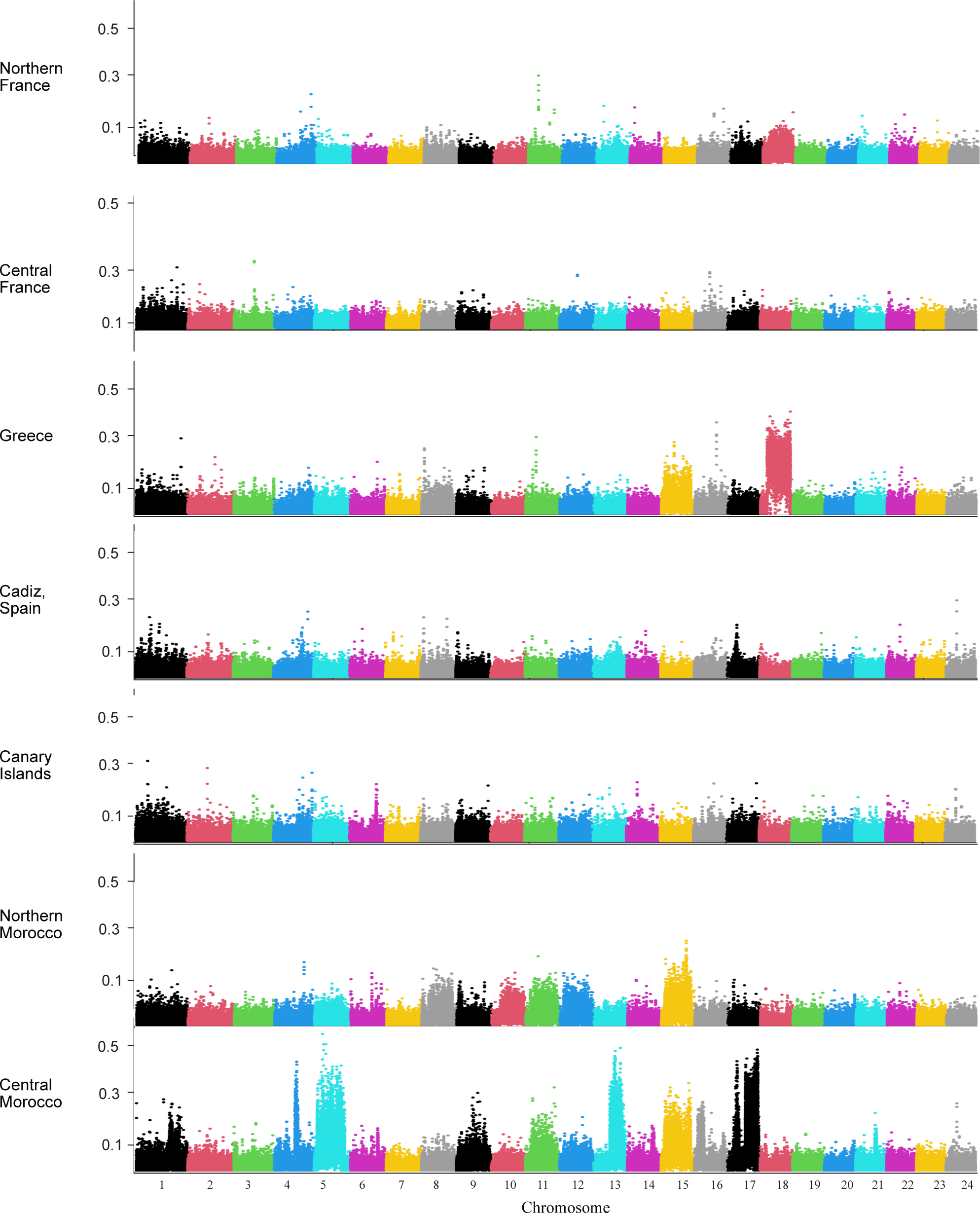
Pairwise F_ST_ values for 20-SNP genomic windows for adult and juvenile populations from the same location. Population codes (see Table 1) for the adult locations are as follows are as follows: NFR (North France), CFR (Central France), AEG (Greece), CAD (Cadiz, Spain), CAN (Canary Islands), NMO (Northern Morocco), and CMO (Central Morocco) (see Figure 1 for more details).

**Figure S5.**
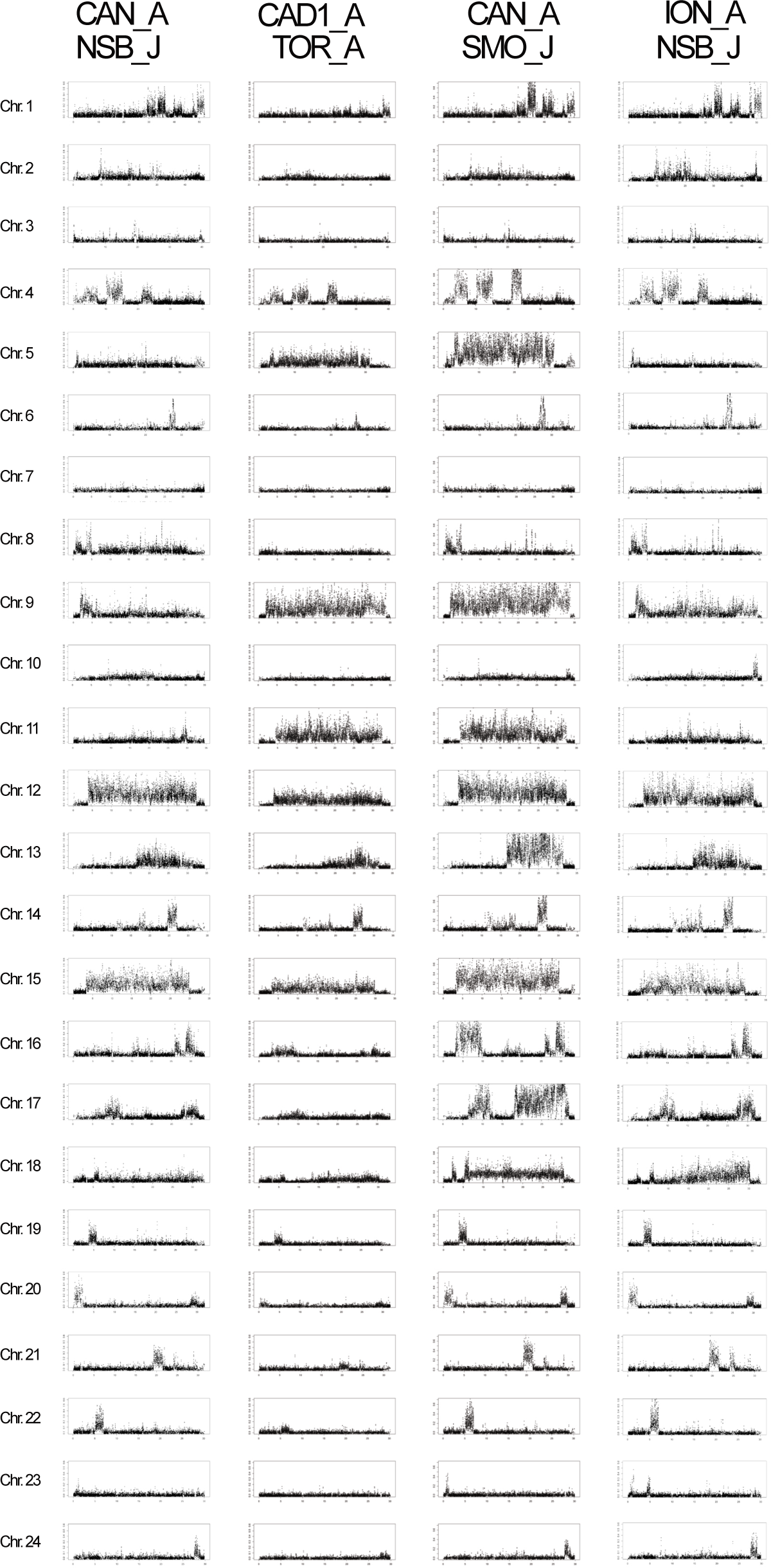
Pairwise F_ST_ values for 20-SNP genomic windows for selected population pairs exhibiting atypical patterns of population differentiation. Population codes correspond to the samples detailed in Table 1: SMO (Southern Morocco), CAN (Canary Islands), NSB (Northern Spain), ION (Ionian Sea), and TOR (Western Mediterranean).

**Figure S6.**
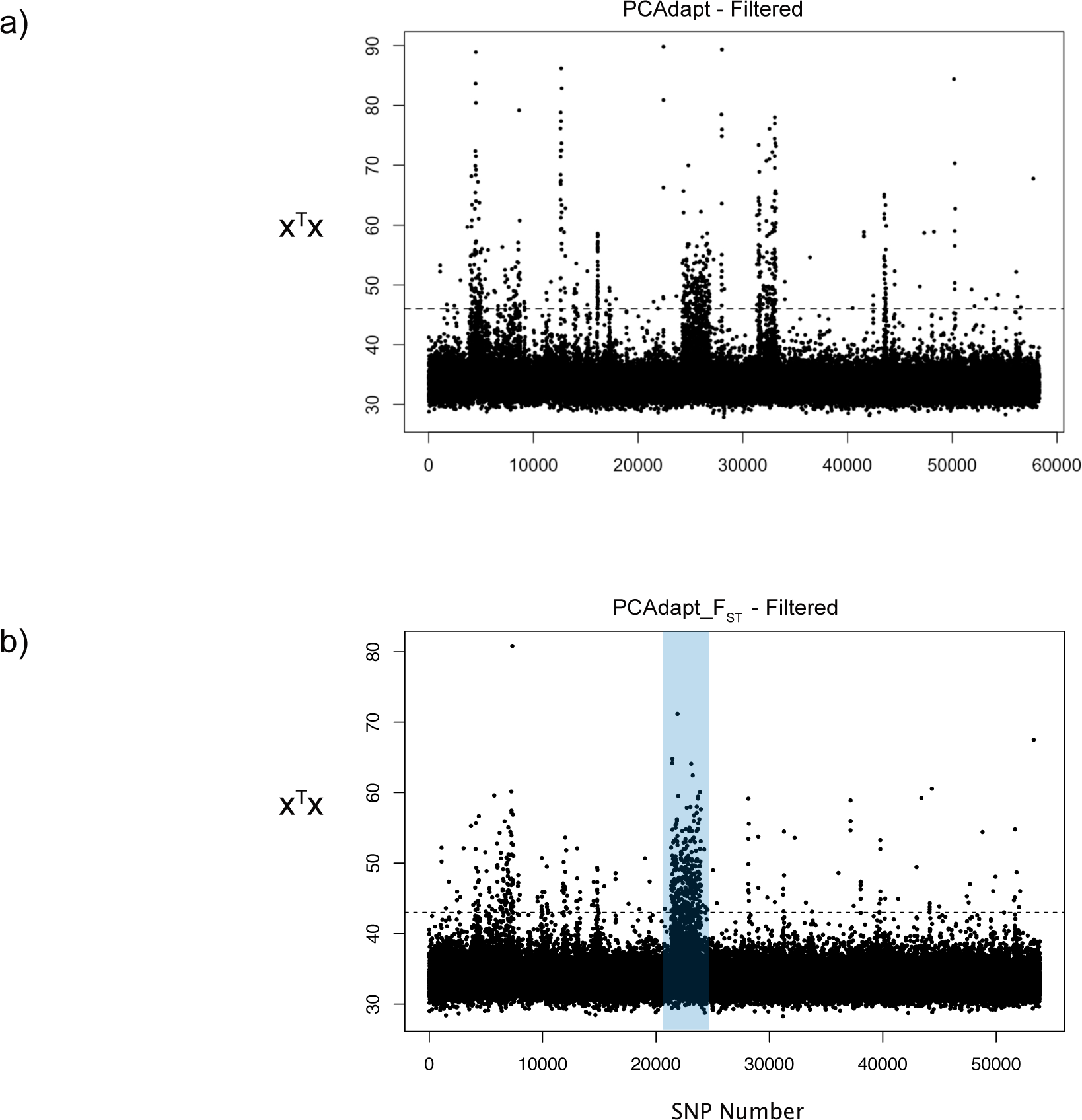
Manhattan plots of xTx (an analog of F_ST_) from the BayPass outlier analysis for the (a) PCAdapt-Filtered datasets and (b) PCAdapt-F_ST_ Filtered datasets. The dashed line in each plot represents the 99th percentile threshold, above which SNP loci were identified as outliers and filtered. In the final dataset, the region shaded in purple was also hard-masked in the “Hard-Filtered” SNP dataset.

**Figure S7.**
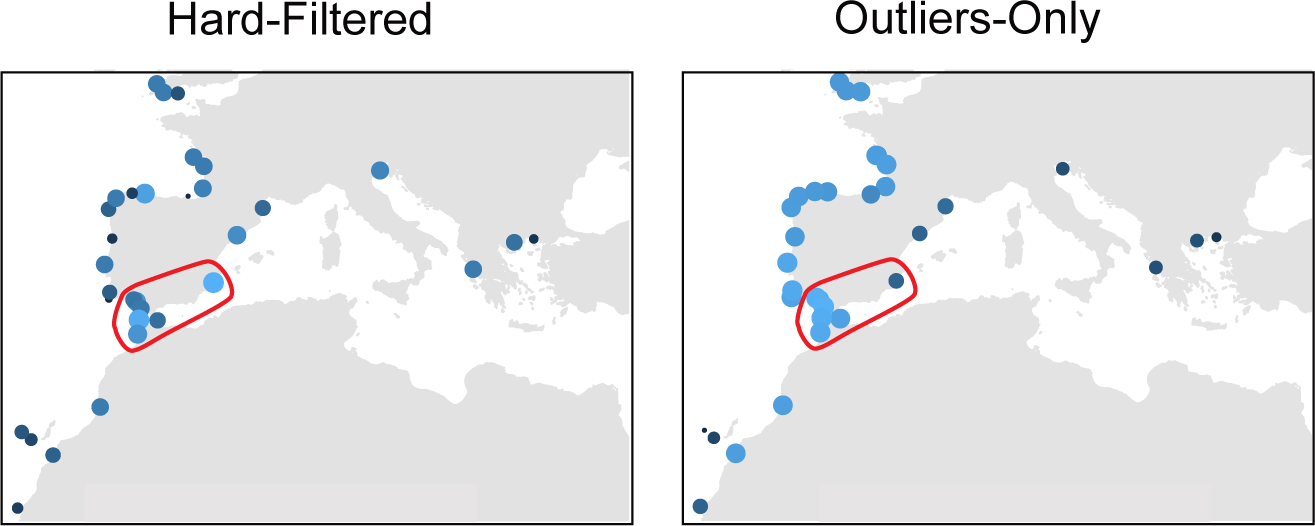
Estimated expected heterozygosity (H_E_) per population of European sardine (Sardina pilchardus), calculated using neutral loci (hard-filtered dataset) and loci within inversions or under selection (outliers-only dataset). The blue circles represent the 34 populations studied, corresponding to the locations listed in Table 1. Circle size and lighter coloration indicate higher H_E_ values. Populations outlined in red consistently exhibited high H_E_ across all filtered datasets.

**Figure S8.**
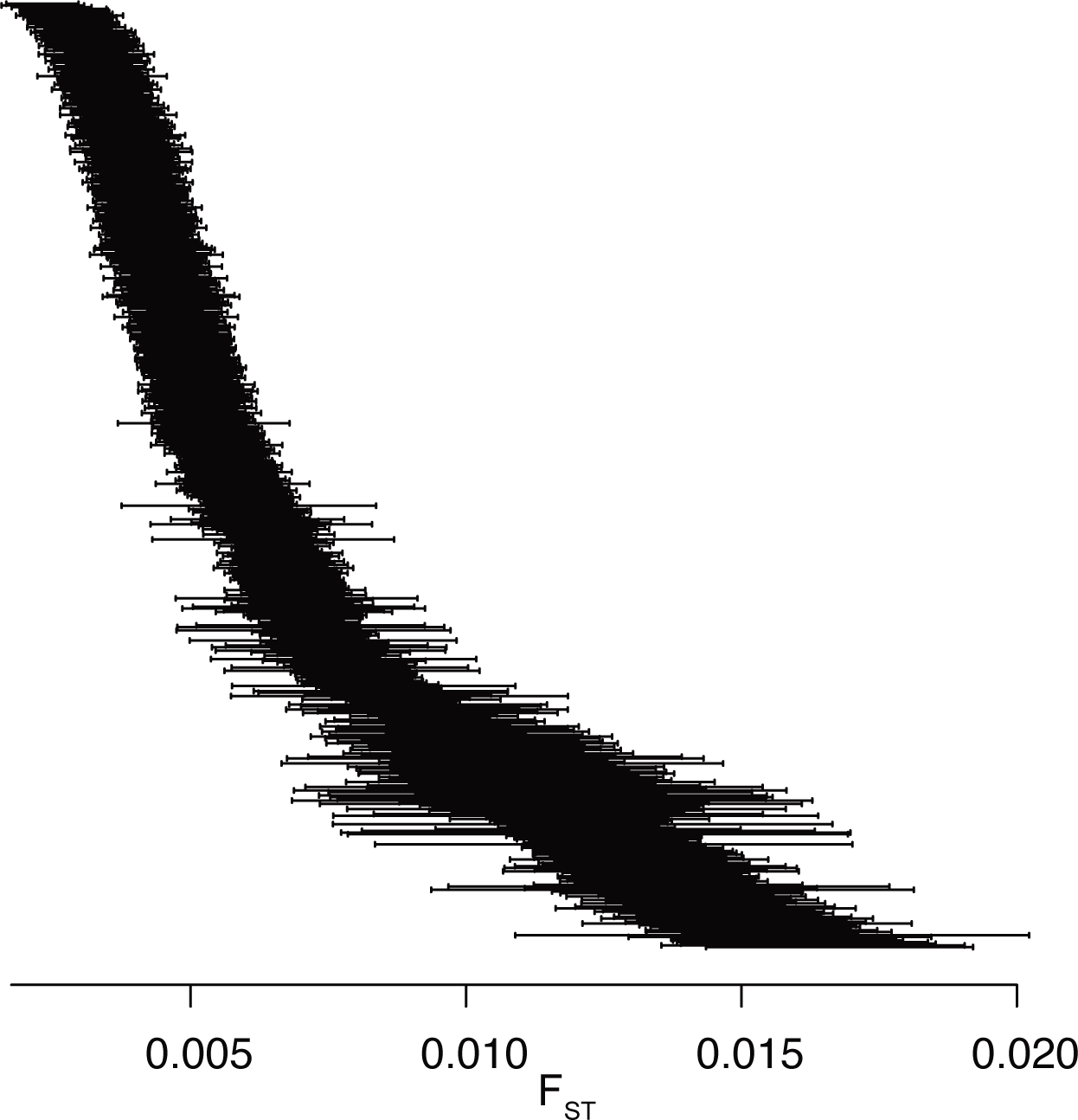
Confidence intervals (95%) for pairwise F_ST_ estimates based on the Hard-Filtered dataset. Values were calculated using the block-jackknife method (see Table S1 for corresponding F_ST_ estimates).

**Figure S9.**
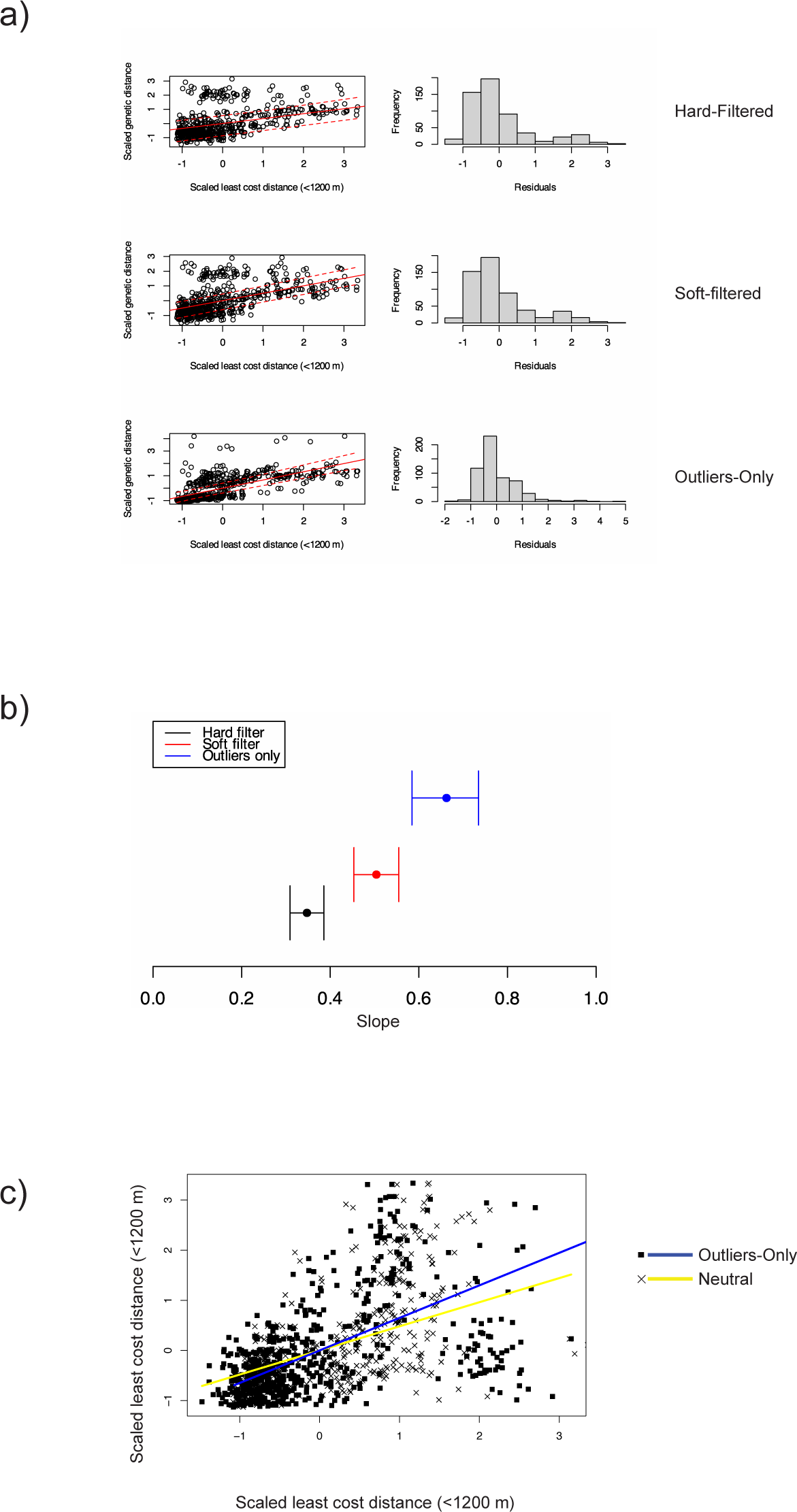
a) Isolation by Distance (IBD) plots for three SNP datasets: “Soft-Filtered” and “Hard-Filtered” datasets, which were filtered to varying degrees for outliers and inversions, and the “Outliers-Only” dataset, which included SNPs from outlier regions. A positive correlation between geographic and genetic distance was observed in all three datasets. (b) Confidence intervals (95%) for slope estimates in each IBD plots in (a). (c) Results of the ANCOVA analysis comparing IBD patterns between the “Outliers-Only” dataset and neutral datasets.

**Figure S10.**
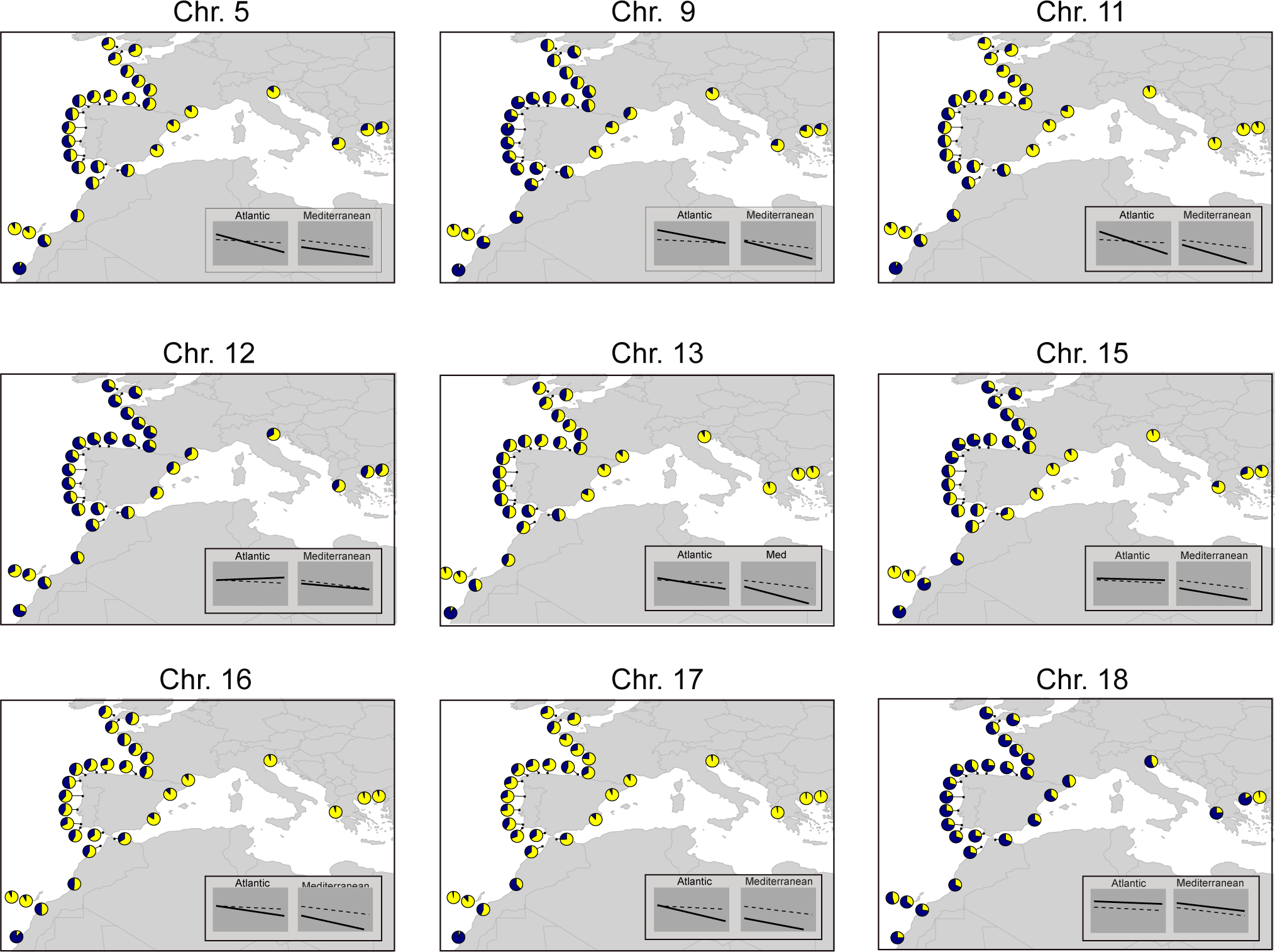
Estimated frequencies of alternative alleles (navy segments in pie charts) for chromosome-scale inversions identified in European sardine populations. Six inversions (Chr. 5, 9, 11, 13, 15, 16, and 17) exhibit a shared pattern of relatively low alternative allele frequencies in the Canary Islands and Mediterranean populations compared to other regions. In contrast, three inversions (Chr. 12, 14, and 18) show alternative allele frequencies in the Mediterranean similar to those in the Atlantic populations. The inversion on Chr. 18, demonstrates how neighboring populations, such as those in Greece, can have significantly different inversion allele frequencies, despite otherwise high levels of admixture. Subplots within each map display trendlines comparing latitudinal (Atlantic) or longitudinal (Mediterranean) gradients in allele frequencies: dotted lines represent regressions for SNPs in non-inverted or outlier regions, while solid lines show those for each inversion. Three population samples (CAD1_J, CAD3_J, and NMO_A), which were adequately represented by other samples, were excluded from the figure to enhance clarity and facilitate the visualization of spatial trends in allele frequencies.

**Figure S11.**
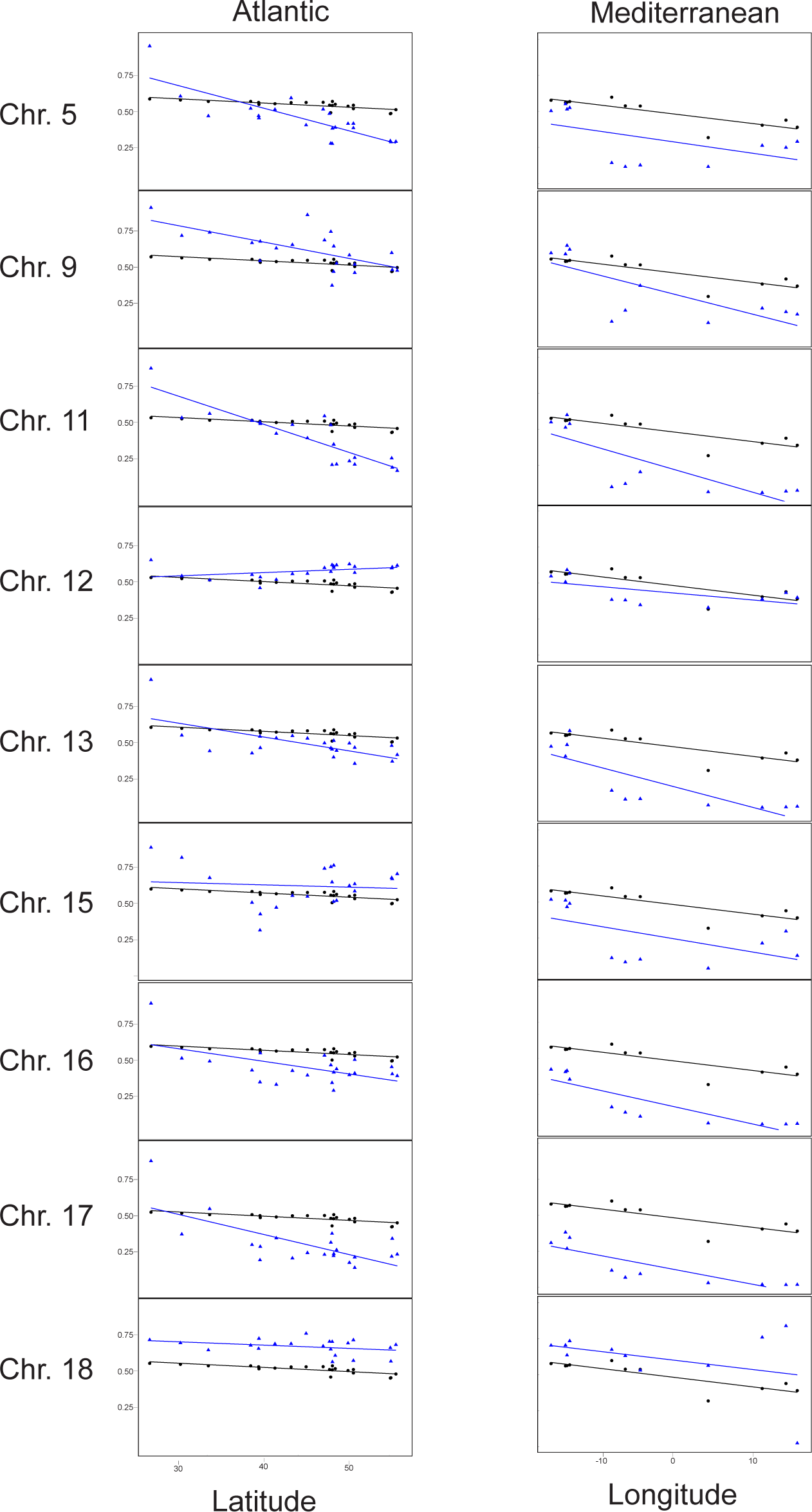
Latitudinal and longitudinal gradients in the frequency of alternative alleles for major inversions in the European sardine. Parametric ANCOVA revealed that the slopes representing allele frequency changes with latitude in the Atlantic Ocean and longitude in the Mediterranean Sea were significantly different (p<0.05). Allele frequencies for SNPs in outlier regions and inversions are shown by blue datapoints and trendlines while those for neutral SNPs are in black.

**Table S1.** Pairwise F_ST_ values between the 34 European sardine populations analyzed in this study. Results are presented for two datasets: the ‘Hard-Filtered’ dataset, which fully masks loci within outlier regions associated with inversions and selection, and the ‘Soft-Filtered’ dataset, which filters only the most extreme loci within those regions (see Methods for filtering criteria). For comparison, results from SNPs within outlier regions using the ‘Outliers-Only’ dataset are also provided. See Supplemental Table S1

**Table S2.**
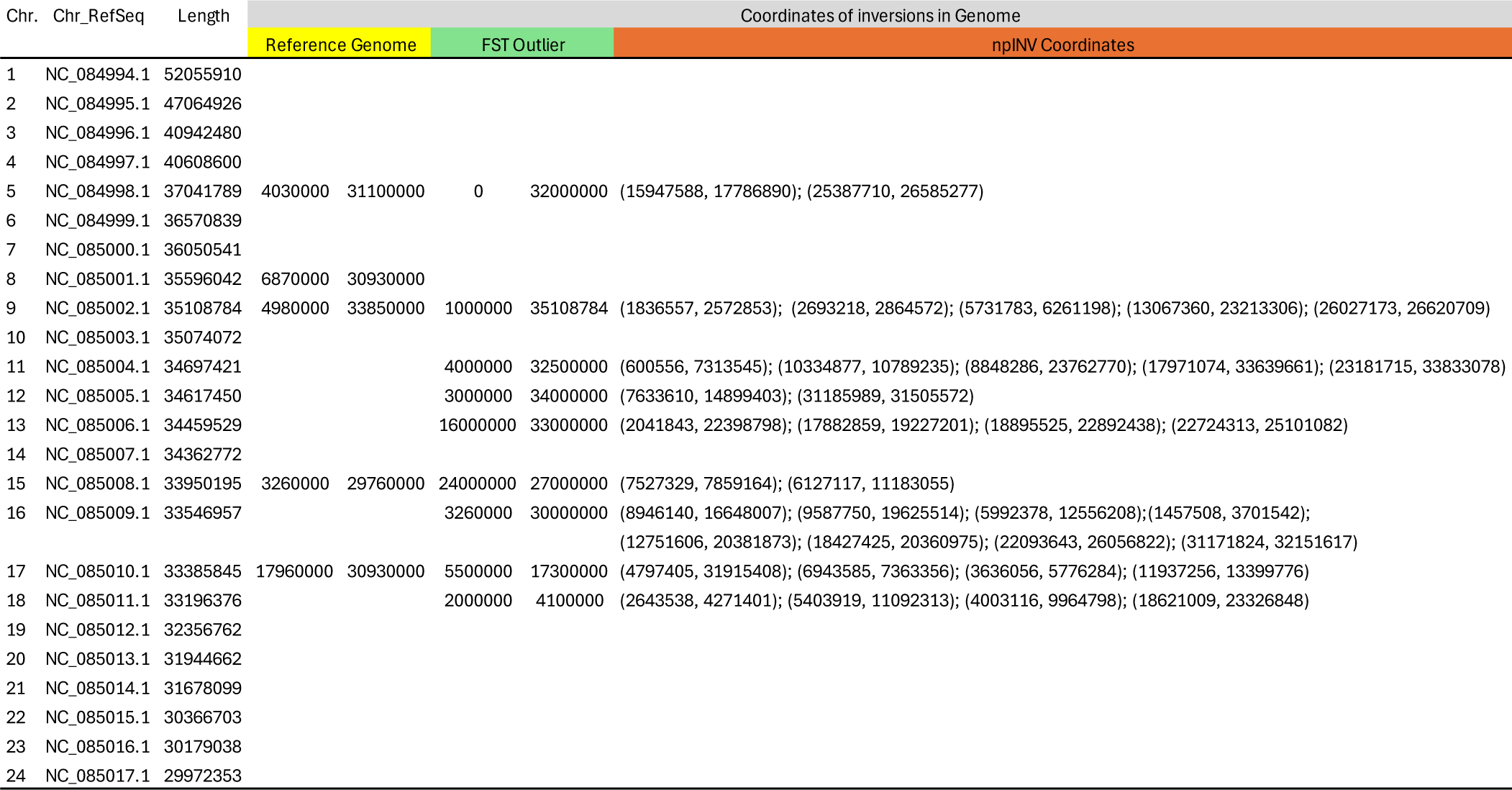
Genomic coordinates of inversions identified in our study using F_ST_-outlier tests and long-read DNA sequencing, compared to those found in the NCBI reference genome.

**Table S3.** Expected heterozygosity (H_E_) estimates for the 34 European Sardine populations studied based on pooled sequencing data. Results are shown for two datasets: the “Hard-Filtered” dataset, which excludes outlier loci associated with inversions and regions under selection (see Methods for filtering criteria), and the “Outliers-Only” dataset, which includes SNPs from loci likely influenced by inversions and selection. Numbers and Codes correspond to sampling locations listed in Table 1.

| Map Num. | Sampling area | Code | Basin | $H_E$ | | |
| --- | --- | --- | --- | --- | --- | --- |
|  |  |  |  | Soft-Filtered | Hard-Filtered | Outliers-Only |
| 1 | South Morocco | SMO_J | Atlantic | 0.169 | 0.158 | 0.180 |
| 2 | Central Morocco | CMO_A | Atlantic | 0.180 | 0.159 | 0.203 |
| 3 | Canary Islands | CAN_J | Atlantic | 0.164 | 0.158 | 0.174 |
| 4 | Canary Islands | CAN_A | Atlantic | 0.159 | 0.158 | 0.161 |
| 5 | Central Morocco | CMO_J | Atlantic | 0.180 | 0.159 | 0.205 |
| 6 | North Morocco | NMO_J | Atlantic | 0.182 | 0.160 | 0.205 |
| 7 | North Morocco | NMO_A | Atlantic | 0.182 | 0.160 | 0.206 |
| 8 | Cadiz, Spain | CAD3_A | Atlantic | 0.183 | 0.159 | 0.208 |
| 9 | Cadiz, Spain | CAD1_J | Atlantic | 0.181 | 0.160 | 0.205 |
| 10 | Cadiz, Spain | CAD2_J | Atlantic | 0.183 | 0.159 | 0.208 |
| 11 | South Portugal | SPA_A | Atlantic | 0.181 | 0.157 | 0.206 |
| 12 | South Portugal | SPA_J | Atlantic | 0.182 | 0.158 | 0.207 |
| 13 | Central Portugal | CPL_A | Atlantic | 0.181 | 0.159 | 0.206 |
| 14 | North Portugal | NPO_A | Atlantic | 0.179 | 0.157 | 0.202 |
| 15 | North Spain | NSS_A | Atlantic | 0.180 | 0.159 | 0.203 |
| 16 | North Spain | NSC_A | Atlantic | 0.180 | 0.159 | 0.202 |
| 17 | North Spain | NSB_J | Atlantic | 0.177 | 0.157 | 0.198 |
| 18 | North Spain | NSA_A | Atlantic | 0.179 | 0.160 | 0.201 |
| 19 | North Spain | NSR_A | Atlantic | 0.179 | 0.158 | 0.199 |
| 20 | Central France | CFR_A | Atlantic | 0.179 | 0.159 | 0.201 |
| 21 | Central France | CFR_J | Atlantic | 0.180 | 0.159 | 0.201 |
| 22 | North France | NFR_J | Atlantic | 0.178 | 0.158 | 0.201 |
| 23 | North France | NFR_A | Atlantic | 0.180 | 0.159 | 0.202 |
| 24 | English Channel | ENC_J | Atlantic | 0.179 | 0.158 | 0.203 |
| 25 | English Channel | ENC_A | Atlantic | 0.179 | 0.159 | 0.200 |
| 26 | Bristol Channel | BRC_J | Atlantic | 0.179 | 0.159 | 0.200 |
| 27 | Morocco | MMO_A | Mediterranean | 0.180 | 0.159 | 0.206 |
| 28 | Torre Vieja | TOR_A | Mediterranean | 0.170 | 0.161 | 0.182 |
| 29 | Tarragona | TAR_A | Mediterranean | 0.170 | 0.160 | 0.180 |
| 30 | Lyon Gulf | LYG_A | Mediterranean | 0.171 | 0.159 | 0.184 |
| 31 | Adriatic Sea | ADR_A | Mediterranean | 0.166 | 0.159 | 0.173 |
| 32 | Ionian Sea | ION_A | Mediterranean | 0.166 | 0.159 | 0.175 |
| 33 | Aegean Sea | AEG_J | Mediterranean | 0.166 | 0.159 | 0.175 |
| 34 | Aegean Sea | AEG_A | Mediterranean | 0.161 | 0.157 | 0.164 |

**Table S4.** Results of the Gene Ontology (GO) enrichment analysis based on the European sardine reference genome annotation from NCBI. Enrichment was assessed using topGO v2.38.1. Results are presented for separate analyses of: each of the nine inversions found in European sardines; all nine inversions combined; and the outlier regions identified in our PCAdapt analysis. See Supplemental Table S4

